# FLI1 and FRA1 transcription factors drive the transcriptional regulatory networks characterizing muscle invasive bladder cancer

**DOI:** 10.1101/2021.11.22.468946

**Authors:** Perihan Yagmur Guneri-Sozeri, Gulden Ozden Yilmaz, Asli Kisim, Ece Cakiroglu, Aleyna Eray, Hamdiye Uzuner, Gökhan Karakülah, Devrim Pesen Okvur, Serif Senturk, Serap Erkek-Ozhan

## Abstract

Bladder cancer is mostly present in the form of urothelium carcinoma, causing over 150,000 deaths each year. Its histopathological classification as muscle invasive (MIBC) and non-muscle invasive (NMIBC) is the most prominent aspect, affecting the prognosis and progression of this disease. In this study, we defined the active regulatory landscape of MIBC and NMIBC cell lines using H3K27ac ChIP-seq and used an integrative approach to combine our findings with existing data. Our analysis revealed FRA1 and FLI1 as two critical transcription factors differentially regulating MIBC regulatory landscape. We show that FRA1 and FLI1 regulate the genes involved in epithelial cell migration and cell junction organization. Knock-down of FRA1 and FLI1 in MIBC revealed the downregulation of several EMT-related genes such as *MAP4K4* and *FLOT1*. Further, ChIP-SICAP performed for FRA1 and FLI1 enabled us to infer chromatin binding partners of these transcription factors and link this information with their target genes. Finally, for the first time we show that knock-down of FRA1 and FLI1 result in significant reduction of invasion capacity of MIBC cells towards muscle microenvironment using IC-CHIP assays. Our results collectively highlight the role of these transcription factors in selection and design of targeted options for treatment of MIBC.

## Introduction

Bladder cancer arising with 90% frequency from urothelium affected more than 500,000 people in 2020 ^1^. According to its histopathological state, bladder cancer is classified as non-muscle invasive bladder cancer (NMIBC) and muscle invasive bladder cancer (MIBC). NMIBC constitutes the great majority of the bladder cancers (80%) and consists of the tumors with the stages Ta and T1 while the remaining MIBC includes the tumors with the stages T2-T4 ^2^. TURBT and BCG options are among the first line treatment choices for NMIBC. On the other hand, MIBC cases are required to have radical cystectomy, chemotherapy and radiotherapy ^3, 4^. Although NMIBC has a better prognosis compared to MIBC, NMIBC patients need a costly long-term follow-up and can develop into MIBC with 20 % frequency ^5^. Recent studies, mainly driven by the TCGA consortium, have characterized the mutational landscape and annotated molecular subgroups of both NMIBC ^6, 7^ and MIBC ^8, 9, 10^. One major finding from these molecular studies was the exceptionally high rate of chromatin modifier mutations both in NMIBC and MIBC. Almost 80% of the bladder cancer patients have mutations in the genes involved in chromatin regulation ^11^. *KMT2D* (28%), *KDM6A* (26%) and *ARID1A* (25%) are the top three highly mutated chromatin modifier genes in bladder cancer ^8^. These findings definitely point out to the epigenetic deregulation and defects in regulation of the gene expression in this disease. Thus, a molecular understanding of the chromatin level regulation is essential. During the last decade, mainly the studies driven by ENCODE and Roadmap consortiums revealed the chromatin and regulatory landscapes of diverse normal and cancer cell lines, and normal tissues ^12, 13, 14, 15^, providing important clues about the regulatory networks characterizing these cell lines and tissues. Identification of the regulatory landscapes of tumorigenic samples has been started to get great attention, given the potential applicability of the knowledge arising with these kind of studies. Especially, characterization of the active enhancer elements in pediatric brain tumors has proven very useful, resulting in identification of the transcription factors and cell of origin differentially involved in molecular subtypes of these tumors ^16, 17^.

There are several studies which analyzed the expression pattern of certain genes in NMIBC and MIBC, which are potentially involved in proliferative and invasive properties of bladder cancer cell lines. *STAG2* expression has been identified as a prognostic marker for NMIBC progression ^18^. In addition, one study suggested a higher rate of progression of T1 stage NMIBC patients with high *CDKN2A* expression and low *FGFR3* expression ^19^. Another study implicated the prognostic value of high *ERBB2* expression in progression of NMIBC ^20^. It was also shown that determining the expression patterns of *FASN, Her2/neu*, and *E2F1* might be important for the correct treatment of NMIBC cases ^21^. High expression of *TGFBI* has been determined to be involved in proliferative and invasive characteristics of bladder cancer cells ^22^. However, there is not yet a study uncovering the regulatory networks differentially characterizing NMIBC and MIBC, which could explain the global molecular mechanisms involved in muscle invasive status of bladder cancer. In this study, we define the active enhancer landscapes of NMIBC and MIBC using H3K27ac ChIP-seq we generated in two NMIBC cell lines, RT4 and RT112 ^23^ and four MIBC cell lines, T24, J82, HT1376 and 5637 ^24, 25, 26^. H3K27ac is a definite marker of active regulatory landscapes previously used in many other studies ^27^. We determined the differentially regulated enhancers between NMIBC and MIBC, integrated the ChIP-seq data with RNA-seq, known gene-enhancer target information and chromatin proteomic assays, and validated our results by functional assays. Our results show that FRA1 and FLI1 are the two transcription factors mainly governing the transcriptional regulatory network of MIBC in interaction with SWI/SNF remodeling complex, and their depletion downregulates the genes involved in epithelial cell migration and decreases the invasion ability of MIBC cell lines.

## Methods

### Cell Lines

T24, J82, HT1376, 5637, RT4 and RT112 bladder cancer cell lines were used in this study. J82 and RT4 cell lines were kindly provided by Şerif Şentürk and T24 cell line was provided by Neşe Atabey. Other cell lines HT1376, RT112 and 5637 were purchased from DSMZ (German Collection of Microorganisms and Cell Lines). T24, J82, RT4 and HT1376 cell lines were grown with Dulbecco’s modified Eagle medium (Gibco) supplemented with 10% fetal bovine serum and 1% penicillin. Other cell lines, RT112 and 5637 were grown in Roswell Park Memorial Institute (RPMI) 1640 Medium (Gibco) supplemented with 10% fetal bovine serum and 1% penicillin. C2C12 myoblast cells were kindly provided by Mehmet Öztürk. Muscle differentiation of C2C12 cells was achieved using 10% horse serum (Gibco, 26050070) for 5 days.

### Chromatin Immunoprecipitation (ChIP)

Chromatin immunoprecipitation (ChIP) experiments were performed with nearly 15 million cells per ChIP. Modified version of the protocol from Weber et al., 2007 ^28^ was followed in all experiments that were performed for each cell line separately. For H3K27ac ChIPs, cells were cross-linked with 1% formaldehyde for 10 min at room temperature, followed by 2.5 M Glycine treatment at +4 °C for 10 minutes to stop the crosslinking. Then, cells were washed and lysed with Lysis Buffer in the presence of 1X Protease Inhibitor Cocktail (Roche,11873580001). Chromatin was fragmented into small pieces (200-500 bp) using S220 Covaris Ultrasound Sonicator. After sonication of the chromatin samples, 50 µl was taken for input fraction and the rest is used for IP. 4 µg H3K27ac antibody (Active Motif, 39133) was coupled to 40 µl Dynabeads M280 Anti Rabbit Magnetic beads (Thermo Fisher, 11204D) (pre-washed with BSA) via rotating overnight at +4°C. Next day, chromatin was bound with pre-washed magnetic beads coupled with H3K27ac antibody for 3 hours. Then, stringent washes were done using a freshly prepared DOC buffer, Lysis buffer and TE. Immunoprecipitated chromatin was eluted by resuspending the beads in a freshly prepared 100 µl elution buffer by agitating the sample at 1000 rpm at 25 °C for 15 minutes, centrifuging the sample at 11000 rpm for 2 minutes at room temperature. Elution step was repeated once more. For transcription factor ChIPs, a slightly more stringent protocol was applied. Different from the H3K27ac ChIPs, cells were cross-linked with 1.25% formaldehyde at RT for 15 minutes and magnetic bead blocking was done using 60 µl magnetic beads in the presence of 10 µl tRNA (10 mg/mL) and 10 µl BSA (10 mg/mL), rotating for 2 hours at +4 °C. Chromatin was precleared by addition of 10 µl pre-blocked magnetic beads and rotating the sample at +4 °C for 1 hour. After chromatin pre-clearing, antibodies targeting transcription factors were added to the chromatin sample in determined concentrations (see Antibodies part of methods) and the sample was rotated overnight at +4 °C. Following day, antibody-bound chromatin was coupled with 50 µl pre-blocked magnetic beads by rotating at +4 °C for 3 hours. All other steps were performed in the same manner as in H3K27ac ChIPs.

For elution of DNA from the input and IP fractions, sample volumes were brought to 200 µl by addition of TE buffer, and 4 µl RNAse (10mg/mL) was added and the sample was incubated for 30 minutes at 37 °C. Then, 1% SDS, 100 mM NaCl and 200 µg/ml Proteinase K were added at final concentrations and the sample was incubated at 55 °C for at least 2.5 hours then overnight at 65 °C. Zymo DNA clean-up & concentrator kit was used to purify the DNA.

### ChIP-SICAP

ChIP-SICAP protocol developed by Rafiee et al., 2016 was utilized ^29^. For the ChIP part of the protocol, T24 cell line was cross-linked with 1.5% final concentration of formaldehyde for 15 min and followed by addition of 150 mM Glycine for 5 minutes, continuously shaking at RT. After a couple of washing and lysis steps, protein concentrations were measured in the presence of 1 % PBS-SDS. Standard proteins were treated the same except Benzonase addition. Then samples were incubated at 95 °C for 5 minutes. Then T24 chromatin’s DNA and proteins were separated by adding 0.5 µl Benzonase (Sigma Aldrich, E8263-5KU) and vortexing. 25 µl from prepared samples were added to one well of 96 well plate and Pierce™ BCA Protein Assay Kit (Thermo Fisher Scientific, 23225) manual was followed to measure the protein content. Overall, for each replicate, chromatin samples corresponding to nearly 2 µg protein were used for ChIP assays performed for FRA1 and FLI1 transcription factors. Each chromatin sample was sonicated for 19 cycles, 1 minute on/1 minute off on Covaris S220. Input DNA of each sample was controlled for sonication size. After sonication, the samples were centrifuged for 10 minutes at 12000g at +4 °C. TritonX-100 was added to the supernatants with a final concentration of 1.5%. Then FLI1 (Abcam, #ab15289) (1:50) and FRA1 (Cell Signaling Technology, #5281) (1:50) antibodies were added to the samples and antibodies were coupled with chromatin samples overnight at +4 cold room with 800 rpm agitation on Thermomixer. Chromatin samples not bound with antibody were used as negative controls. Next day, the samples were centrifuged at +4 °C for 10 minutes at 12000 g. Supernatants were taken and filled up to 1 mL with IP Buffer ^51^. Meanwhile, 50 µl Dynabeads (M280 Anti Rabbit Magnetic Beads) for each sample were washed with IP Buffer (+4 °C) once. Pre-washed magnetic beads were then added to the samples and coupled at +4 °C for 3 hours by rotating. Then samples were washed and resuspended with 200 µl PBS-T (PBS including 0.1% Tween20) and shipped to Francis Crick Institute for the SICAP part of the protocol. After the chromatin immunoprecipitation, the chromatin fragments were kept bound to the protein A/G beads on dry ice until they were treated using the ChIP-SICAP procedure. ChIP-SICAP was performed as already described in ^51, 95^. Briefly, the beads were treated with Klenow 3’-exo minus, T4 PNK, and dNTPs (NEB) to make 3’-overhangs and remove 3’-phosphates. Then, the beads were treated with terminal deoxynucleotidyl transferase (TdT) and biotinylated nucleotides (ddUTP and dCTP 1:1). The beads were then washed 6 times with IP buffer (Tris-HCl pH 7.5 50 mM, Triton X-100 1%, NP-40 0.5%, EDTA 5 mM). The isolated proteins were eluted using elution buffer (SDS 7.5%, DTT 200 mM) by incubating at 37 °C for 15 min. Eluted samples were diluted in IP buffer. Then 100 μL of either prS beads were added for the DNA enrichment. Streptavidin beads were washed 3 times with SDS washing buffer (Tris-HCl 10 mM pH 8, SDS 1%, NaCl 200 mM, EDTA 1mM), once with BW2x buffer (Tris-HCl pH 8 10 mM, Triton X-100 0.1%, NaCl 2M, EDTA 1 mM), once with iso-propanol 20% in water and three times with acetonitrile 40% in water. The beads were transferred to PCR tubes using acetonitrile 40%. The beads separated on the magnet, and the supernatant was removed. Then the beads were resuspended in 15 μl Ambic 50 mM plus DTT 10 mM final concentration. Then the samples were incubated at 50 °C for 15 min to reduce the disulfide bonds. The cysteines were then alkylated with IAA 20 mM final concentration for 15 min in dark. IAA was neutralized by adding DTT 10 mM final concentration. To digest the proteins, 300 ng LysC (Wako) was added to each sample. After an overnight incubation, the supernatant was transferred to a new PCR tube. Then 200 ng Trypsin was added to each tube. The digestion continued for 6-8 hours. Finally, the peptides were cleaned up using Ziptips with 0.6 µl C_18_resin (Merck). The samples were injected to an Orbitrap Fusion Lumos, working on data dependent acquisition mode.

### Antibodies

ChIP experiments were carried out with 5 µg H3K27ac antibody (Active Motif, #39133), 1:100 diluted FRA1 antibody (Cell Signaling Technology, #5281) and 1:100 diluted FLI1 antibody (Abcam, #ab15289, Cell Signaling Technology #35980). For ChIP-SICAP, FLI1 and FRA1 antibodies were used with 1:50 dilution. For Western Blotting, FRA1 and FLI1 antibodies were used with 1:1000 dilution. B-actin primary antibody (Cell Signaling Technology, 3700) was used as reference protein with a 1:1000 dilution. Anti-Rabbit Secondary Antibody in 1:30000 dilution and Anti-Mouse Secondary Antibody (Cell Signaling Technology, 5151; Li-Cor, 926-68020, respectively) were used in 1:15000 dilution during Western Blotting experiments.

### Western Blotting

48 hours and 72 hours after knockdown, cells were collected with 200 µl RIPA + PIC (1X) Buffer. Samples were vigorously vortexed in every 10 minutes up to 30 minutes. Then the samples were sonicated for 5 cycles (30 seconds on/ 30 seconds off) to efficiently get the proteins in the nucleus, too. After sonication, the samples centrifuged for 20 minutes at +4 °C at full speed. We stored the supernatant of the samples at −80 °C. Protein amounts were measured according to the Pierce™ BCA Protein Assay Kit (Thermo Fisher Scientific, 23225)’s protocol. Then, the proteins were diluted with Laemmli Buffer and RIPA Buffer (1X) to equal protein concentrations of the samples and boiled at 95 °C for 5 minutes. 30 µg proteins were loaded into %8 separating gel. Bio-RAD Western Blot System (1658029) was used to perform Western Blotting. Samples were first run at 90 Volt (V) for 20 minutes until the proteins passed into separating gel and then ran at 120 V for 1 and a half hours. Samples were then transferred to nitrocellulose membrane at 350 mA for 1.5 hours. After transferring the proteins to the membrane, membrane was blocked with 5% milk powder dissolved in TBS-T for 1 hour on a shaker at RT, followed by the antibody binding overnight at +4 °C on a shaker. The next day membranes were washed with TBS-T for 10 minutes 3 times on a shaker at RT. Then, secondary antibody was bound to primary antibody on the membrane for 1 hour at RT on a shaker, which was followed by washing with TBS-T three times again. Membranes were imaged using Li-COR ODYSSEY Clx device.

### RT-qPCR

RT-qPCR experiments were performed to analyze the changes in gene expressions after control siRNA, FLI1, FRA1 knockdown or FLI1 and FRA1 knockdown together. 48 hours and 72 hours post-transfection of siRNAs, RNA from the cells were collected using MN Nucleospin RNA Isolation Kit (740955.50) according to kit’s instructions. Elution volume was 40 µl for each sample. Collection of RNA was followed by cDNA conversion using 1 µg RNA for each reaction with the Maxima First Strand cDNA Synthesis Kit (Thermo Scientific, K1642). Expression of the genes were determined using FastStart Essential DNA Green Master Kit (ROCHE, catalog no: 06402712001). cDNA converted from RNA was used as 1:5 dilution (in ultra-pure water) per reaction. Quantitative PCR reaction was held on Applied Biosystems 7500 Fast Real Time PCR Device according to the manufacturer’s instructions. Relative expression of the target genes were calculated using the 2^ΔΔCT Method ^30^. For each biological replicate, RT-qPCR experiments were done in three technical replicates and mean expression values for the two biological replicates values were plotted. Plots representing relative expression changes were drawn at GRAPHPAD version 8.3.0 ^31^.

### Primer Design

All primers were designed using NCBI Primer Blast Tool ^32^. Sequences of the primers are listed in Supplementary Table 7.

### Knockdown Experiments

Custom siRNAs targeting 5’-CACCAUGAGUGGCAGUCAG-3’ ^33^ for FRA1 and 5′-GUUCACUGCUGGCCUAUAA-3’ ^34^ for FLI1 were ordered from GE Healthcare Dharmacon, Inc. As negative control, MISSION^®^ siRNA Universal Negative Control SIC001 was used (MERCK). In each knockdown assay, siRNAs were transfected using Thermo Fisher Lipofectamine 3000 Transfection Reagent (Catalog Number: L3000001) according to transfection reagent’s protocol. 75 nM siRNA was transfected to T24 cells (65% confluent) for FLI1 and FLI1 unique knockdowns and 50 nM siRNAs targeting FLI1 and FRA1 were used for co-knockdown of FLI1 and FRA1. T24 cells were incubated in transfection medium for 24 hours. Then, the transfection medium was removed and transfected cells were passaged further to be collected for RNA and protein isolation.

### Invasion assay - IC-Chip Design and Analysis

We used IC-CHIP (INITIO Cell Biyoteknoloji) to test the muscle invasion capacity of T24 and 5637 cell lines for control siRNA, FLI1 siRNA, FRA1 siRNA and co-knockdown of FLI1 and FRA1. Differentiated C2C12 myoblast cell line was used to mimic “muscle microenvironment”. C2C12 cell line differentiation condition was set up by supplementing the C2C12 cells with 10% Horse Serum (Sigma Aldrich, H1138-100ML) rather than FBS for 5 days. One day before setting up the invasion assays, T24 and 5637 cells were labeled using 5 µM Green CellTracker (Thermo Fisher) according to manufacturer’s instructions. Knockdown experiments were done as mentioned above. Growth Factor Reduced Matrigel (Corning, 354230) (1:1 diluted with FBS free medium) was added to the middle channel of the IC-CHIP to set positive and negative controls’ middle channel environment and Growth Factor Reduced Matrigel (1:1 diluted with differentiated C2C12 cells at 1 million cells/ 1 mL in FBS free medium) was added to the middle channel to monitor “invasion to muscle” (Please see Fig. 6a for the setup). Afterwards, IC-CHIPs were incubated at 37 ° C for 30 minutes. Then bottom channels of the IC-CHIPs for positive and negative controls were loaded with FBS(+) medium and FBS(-) medium, respectively. Upper channels were loaded with T24 or 5637 cells at 1 million/cells in FBS(-) medium for positive and negative control conditions. For the “invasion to muscle” setup, T24 or 5637 cells for each condition (control siRNA, FRA1 siRNA, FLI1 siRNA, FLI1 + FRA1 siRNA) were loaded to the upper channel of IC-CHIPs in FBS(-) medium after overnight incubation of the IC-chips containing the muscle microenvironment and FBS(-) medium in upper and lower channels. IC-CHIPs were observed for 3 days (day 0, 1 and 2) post-loading of the T24 or 5637 cells. Images were acquired for each day with a 10X objective using a confocal microscope (Zeiss) and analyzed using Image J ^35^. The quantification of images was performed by Python programming and R Studio as previously described ^36^. The invasion capacity of the cells was determined through normalization of datasets to day 0 ^37^ for T24 and to day 1 for 5637 cells.

### Overexpression of FRA1 and FLI1 in NMIBC cell lines

EF1a_FLI1_P2A_Hygro_Barcode vector (Addgene,120437) was used for FLI1 overexpression and EF1a_mCherry_P2A_Hygro_Barcode vector (Addgene, 120426) was used as control. For FRA1 overexpression, FRA1 sequence from p6599 MSCV-IP N-HAonly FOSL1 vector (Addgene, 34897) was cloned into pLV-EF1α-IRES-puro vector (Addgene, 85132) and pLV-EF1a-IRES-Puro vector was used as control. HEK293T cells were seeded in a 6 well plate and co-transfected with target vector, psPAX2, and pMD2.G at ratio of 4:2:1 using PEI. The supernatant was harvested after 48 and 72 hours and filtered through a 0.45 μm SFCA filter. RT112 cells were infected for 48 hours with either vector and selected with puromycin (2 µg/ml) or hygromycin (200 µg/ml). Antibiotic selection was continued until all the cells in the control were dead.

### Wound Healing Scratch Assay

Wound healing assay was performed to see the effects of overexpression on migration. Cells were seeded in a 6-well plate. When the cells were confluent, the cell layer was scraped from top to bottom in a straight line using a 200 µL tip. The cells were then washed twice with 1x PBS. Cell medium containing 2% fetal bovine serum (FBS) was added to the wells. Images of the cells were taken at 10x magnification at different time points (0h, 6h, 24h, 30h). The results were analysed using ImageJ’s Wound Healing Size tool, with “area inches” value as 0h=100, relative to other time points. Experiment was performed as at least 5 independent measurements

### MTT

Cells were seeded in a 96-well plate as three technical replicates. Cell medium with no-cell wells were used as a blank. After four days, MTT reagent and solubilization solution were added to all wells four hours apart, as recommended by the manufacturer (Cell Proliferation Kit 1, 11465007001 Roche). After overnight incubation, absorbance of wells were measured at 570 nm wavelength.

### Next Generation Sequencing

ChIP library preparation and sequencing were performed at the GeneCore unit of EMBL and Macrogen. In GeneCore, Sequencing libraries were sequenced on the Hiseq2500 platform using single-end 50 bp reads for H3K27ac ChIP and input samples. Transcription factor ChIP and input libraries (FRA1 and FLI1) were sequenced on NextSeq 500 high output mode using 75 bp single end reads and NovaSeq6000 using 150 bp paired end reads.

### Alignment and Processing of NGS Data

Next generation data was aligned to the GRHg38 version of the human genome using Burrows Wheeler Aligner (BWA) version 0.7.17-r1188 with default parameters^38^. Quality of the alignments was confirmed using Homer NGS Data Quality Control Analysis tool version 4.10.3. (makeTagDirectory command) ^39^.

### Gene Annotation

GENCODE Comprehensive Gene Annotation v30 was used to annotate the genes ^40^.

### Peak Finding

Macs2 peak finding algorithm ^41^ was used with -broad option to call the peaks from H3K27ac ChIP-Seq data. Afterwards, the peaks which are completely located in promoter regions (+/-1kb surrounding transcriptional start sites) of protein coding genes (according to the GENCODE v30 Comprehensive Gene Annotation data, https://www.gencodegenes.org/human/release_30.html) were excluded for any further analysis.

FRA1 and FLI1 transcription factor peak finding was performed using 2 biological replicates with Homer getDifferentialPeaksReplicates.pl command, specifying the style as factor (-style factor), adjusting fold enrichment over input tag count to 2 (-F 2), fold enrichment over local tag count 2 (-L 2) and specifying genome to hg38 (-genome hg38) ^39^.

### Finding MIBC/NMIBC Specific Enhancer Regions

DiffBind Bioconductor package with default parameters was used to identify group specific enriched (muscle-invasive bladder cancer specific or non-muscle invasive bladder cancer specific) enhancer ^42, 43^. 4 cell lines in MIBC and 2 cell lines in NMIBC are used as replicates of each group. Using FDR cut-off =< 0.05, we identified 295 NMIBC-specific and 1404 MIBC-specific enhancers.

### Snapshot Visualization of ChIP-seq Signal

Aligned H3K27ac, FLI1 and FRA1 ChIP-seq reads were RPKM normalized using deepTools’ “bamCovarage” function with “normalizeUsing RPKM” parameter ^44^. After normalization, we visualized the ChIP-Seq reads using GViz Bioconductor package ^45^. We used “horizon” type visualization of the reads by adjusting the horizon scale to 50 in Fig. 1, Fig. 2d-f, and Supplementary Fig. 1, 25 in Fig. 4 e-f and Supplementary Fig. 7b, 50 for Fra1 35 for FLI1 snapshot in Supplementary Fig. 9. GENCODE hg38 Comprehensive Gene Annotation Version 30 data was defined as the “GeneRegionTrack” for the annotation of the genes at the region of interest. Transcripts on those regions were shown in collapsed form using “collapseTranscripts” parameter.

**Fig. 1.**
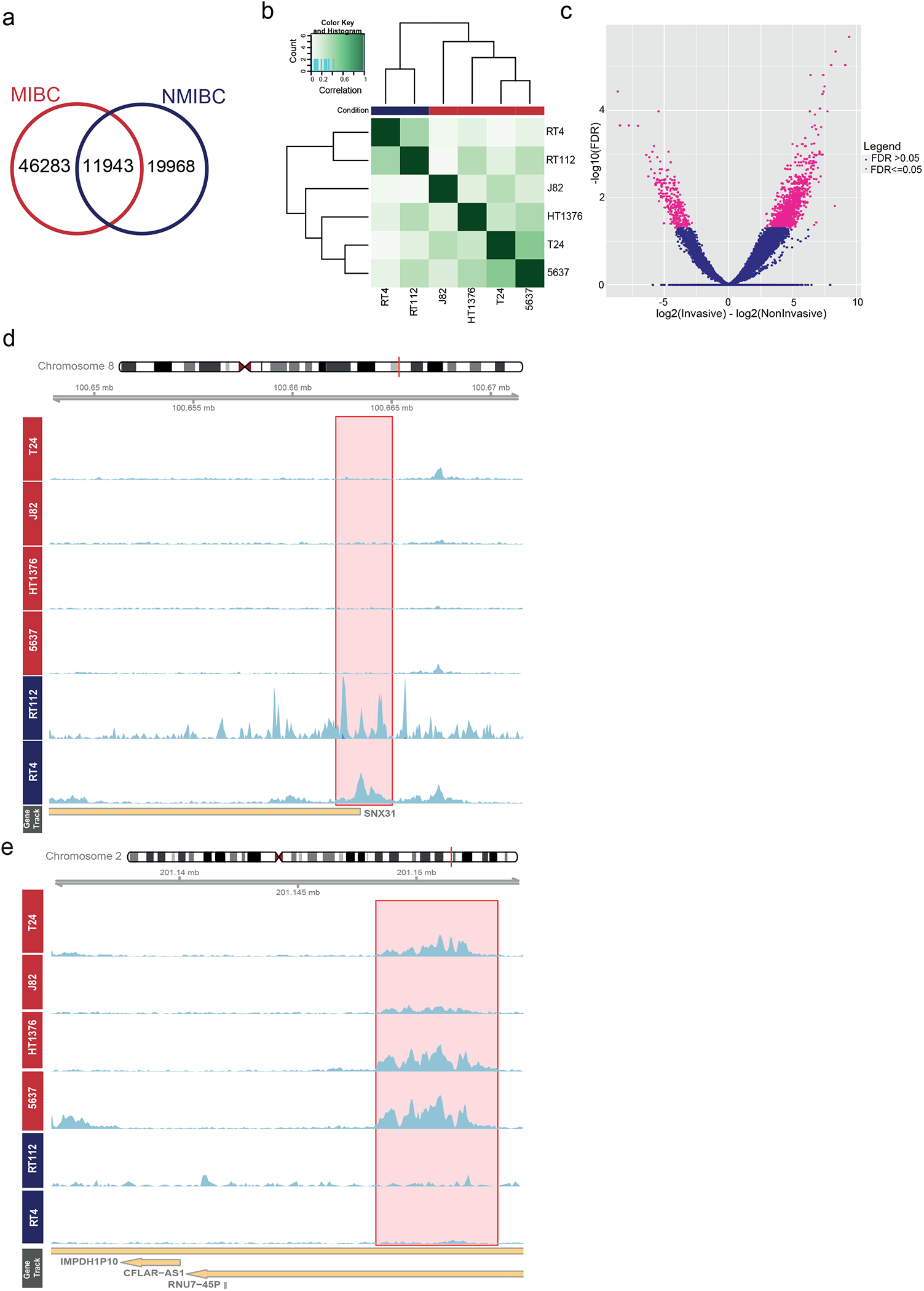
Differentially regulated enhancers between NMIBC and MIBC cell lines. (a) Venn diagram showing the overlap between H3K27ac peaks called in NMIBC and MIBC. (b) Heatmap displaying the correlation among the cell lines according to the signals present at consensus H3K27ac peaks (unsupervised). (c) Volcano plot showing the differentially regulated enhancers (FDR =< 0.05) between MIBC and NMIBC (n=1699). (d,e) Snapshots visualize H3K27ac signal at exemplary NMIBC (d) and MIBC (e) specific enhancers. Scale of the snapshots are adjusted to 50.

**Fig. 2.**
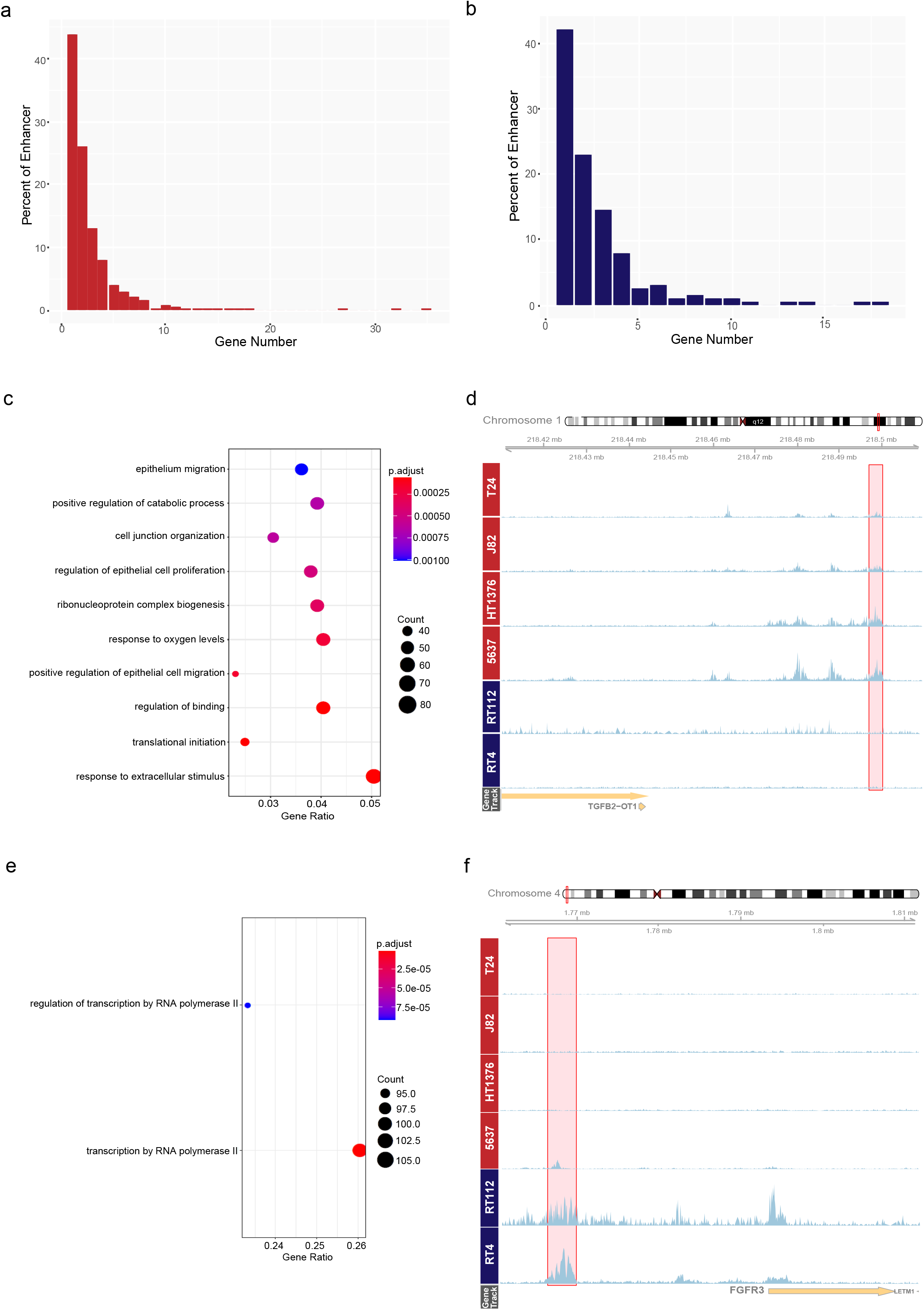
Target genes of MIBC enhancers are involved in regulation of epithelial cell migration. (a, b) Barplots show the relationship between percent of the enhancers and the number of their target genes, (a) MIBC specific enhancers, (b) NMIBC specific enhancers. (c, e) ClusterProfiler outputs displaying the enriched GO terms for the target genes of MIBC enhancers (c) and NMIBC enhancers (e). (d, f) Snapshots showing H3K37ac signal at MIBC enhancer regulating *TGFB2* (d) and NMIBC enhancer regulating *FGFR3* (f). Enhancer regions are shown in pink rectangles. Scale of the snapshots are adjusted to 50.

### Identification of the Expressed Genes in NMIBC and MIBC Cell Lines

We downloaded the gene expression data (rpkm values) from Cancer Cell Line Encyclopedia ^46^ for the NMIBC and MIBC cell lines. To determine a cut-off for the gene expression values, we first calculated the average gene expression values in NMIBC or MIBC and then we used the R package Mclust ^47^ to set a cut-off for gene expression in NMIBC or MIBC cell lines. We took all the gene expression values for protein-coding genes provided in the CCLE database, and removed the genes with expression values ‘0’ rpkm. Then using “G=3” option and p-value <0.05 in Mclust library, we divided the gene expression values (log2(RPKM)) into “expressed” (Group 2 and 3) and “not expressed” (Group 1) (Supplementary Fig. 2). Accordingly, RPKM value of expression cut-off for NMIBC is determined as -0.77 while it is 0.98 for MIBC on log2 scale. These processes resulted in identification of 11681 and 10131 numbers of genes being expressed in NMIBC and MIBC cell lines, respectively.

### Processing of NMIBC and MIBC Patient Gene Expression Data

306 Urothelial Carcinoma samples’ gene expression data was obtained from GSE32894 Accession Code ^48^. Non-normalized gene expression data was normalized by using “normalizeQuantiles” function in Limma Bioconductor Package ^49^. Tumour samples having the stage <T2 were defined as a member of NMIBC group while remaining ones (>=T2) were defined as in MIBC group. Expression boxplots of each group specific transcription factor were drawn by “ggplot2” package function “ggplot” ^50^. To add the significance scores, “ggsignif” package was used by using basic Student’s t test for statistical analysis ^51^.

### Identification of the Genes Regulated by Group Specific Enhancer Regions

Group specific enhancer regions (FDR =< 0.05) were given as a query to UCSC (https://genome.ucsc.edu/cgi-bin/hgTables) geneHancer algorithm to get the predictions of the genes regulated by these enhancers ^52^. GeneHancer Interactions (GH Interactions) table format was chosen as output. Use of the gene-enhancer interaction data provided in GeneHancer database resulted in identification of 3306 and 9269 target genes for NMIBC and MIBC specific enhancers, respectively. However, as all these genes would not be relevant for NMIBC and MIBC cell lines, we filtered out the genes which were not expressed in NMIBC or MIBC and from those kept the protein coding ones. After this filtering step, 456 target genes for NMIBC-specific enhancers and 1752 target genes for MIBC-specific enhancers were identified.

### Transcription Factor Motif Binding Analysis On Respective Regions Including Group Specific Peaks

Tag Directories which were established by using HOMER algorithm’s “makeTagDirectory” command were combined for NMIBC and MIBC separately. Then, nucleosome free regions in these combined tag directories were found using HOMER algorithm’s findPeaks command with -nfr option. Resulting NMIBC or MIBC specific NFRs were overlapped with the respective group specific enhancers. Afterwards, transcription motif finding was performed on the overlapping NFR regions using findMotifsGenome.pl command of Homer algorithm with hg38 and –size given options ^39^.

Motif finding on FLI1 and FRA1’s found united peaks was done using findMotifsGenome.pl command with hg38 and –size 300 options (Supplementary Fig. 8).

### Overlap of MIBC-specific and NMIBC-specific Enhancers with TF Peaks

Overlaps between FLI1 and FRA1 peaks and MIBC or NMIBC-specific enhancers were using -findOverlaps function with -type within parameter using GenomicRanges package ^53^. Output results were visualized as barplots (Fig. 4a).

### Identifying The Target Genes of FRA1 and FLI1 Transcription Factors On MIBC-specific Enhancer Regions

Group-specific enhancer regions targeting a gene (456 target genes for NMIBC-specific enhancers and 1752 target genes for MIBC-specific enhancers) were overlapped with the FRA1 and FLI1 peaks. In this way FLI1 and FRA1 peaks were linked with the target genes. As a result, 557 genes were found to be regulated by FRA1 on MIBC-specific enhancers yet 260 genes were identified as FLI1 targets. Gephi Software (version 0.9.2) ^54^ was used to visualize the transcription factors - target gene networks for the genes involved in epithelium migration, cell junction organization and regulation of epithelial cell proliferation related terms (Fig. 4b).

### Pathway Enrichment Analysis

Pathway enrichment analysis was done using the protein coding ones (according to the GENCODE v30 Comprehensive Gene Annotation data, https://www.gencodegenes.org/human/release_30.html). To analyze the Gene Ontology Terms related with the target genes, they were converted to Entrez gene IDs at first. 1606 genes out of 1752 MIBC target genes and 403 out of 456 NMIBC target genes can be converted to Entrez gene IDs by using mapIDs() function of org.Hs.eg.db package version 3.8.2 ^55^. Final set of target genes of NMIBC was analyzed by using clusterProfiler package (version 3.12.10) enrichGO() function, setting maxGSsize (maximum gene ontology set size) parameter to 3000 and selecting p adjust method as FDR ^56^. On the other hand, the finalized set of the target genes of MIBC group was analyzed by using the same package and function by using default maxGSsize which is 500 and with the same p adjust method. We represented the results of the Gene Ontology term analysis by using the dotplot() function in the clusterProfiler package.

### Analysis of MAP4K4 Expression Data in Primary Tumors

Scatter plots showing the correlation between the expression of MAP4K4 and FRA1 and FLI1 (Supplementary Fig. 11a, b) were generated using the gene expression data available for TCGA 2017 (n=408) cohort at cBioPortal database ^57^ and the tab ‘Plots’. For the metastatic site distribution plots (Supplementary Fig. 11c, d), patients with high MAP4K4 expression were determined using the option ‘mRNA expression z-scores relative to diploid samples (RNA Seq V2 RSEM)’, and resulting clinical data was inspected for the metastatic site distribution. (TCGA 2017 group (n = 408) has MAP4K4 mRNA altered group (n=15) and MAP4K4 mRNA unaltered group (n = 393). On the other hand, TCGA 2014 group has 131 samples, 5 of them is MAP4K4 mRNA altered and the rest (n=126) is unaltered.)

### Analysis of ChIP-SICAP Data

Peptides were injected into Orbitrap Fusion Lumos LC - MS/MS Device working on data dependent acquisition mode. Mass spectrometry raw data were processed with MaxQuant (1.6.2.6) ^58^ using default settings. MSMS spectra were searched against the Uniprot databases (Human Swissprot) combined with a database containing protein sequences of contaminants. Enzyme specificity was set to trypsin/P and LysC, allowing a maximum of two missed cleavages. Cysteine carbamidomethylation was set as fixed modification, and methionine oxidation and protein N-terminal acetylation were used as variable modifications. Global false discovery rate for both protein and peptides were set to 1%. The match-between-runs and re-quantify options were enabled. Intensity-based quantification options (iBAQ) was also calculated. Differentially enriched proteins in comparison to the no-antibody controls were determined using empirical Bayes moderated t-test by limma package ^49^. Multiple-testing corrections were performed using the Benjamini-Hochberg procedure to calculate adjusted p-values. Proteins with adjusted p-value < 0.05 and logFC > 1 were determined as significantly enriched in the respective transcription factor chromatin bound interactome.

Cellular localization of FLI1 and FRA1’s protein partners significantly enriched (logFC > 1, adjusted p value < 0.05) were found using Gene Ontology Terms GO0005634, GO0005694 and GO0033279 ^59, 60^ and plotted using ggVennDiagram and ggplot2 packages ^50, 61^ (Fig. 5a). Scatter plots highlighting some of the important protein interactors were plotted using ggplot2 package ^50^ (Fig. 5b-c).

### Integration of ChIP-SICAP Data with Existing ChIP-seq and DepMap Data

We searched for the ChIP-seq data available for the components of SWI/SNF complex, which are identified to interact with FRA1 and/or FLI1 according to ChIP-SICAP results, in the Cistrome database ^62^. For each factor, we inspected the good quality ChIP-seq data available for epithelial cancer cell lines which were not subjected any special treatment before ChIP experiments at UCSC genome browser ^63^. (please see Supplementary Fig. 13a for the used data sets). MIBC enhancers regulating MAP4K4 and FLOT1 were checked for SWI/SNF complex component ChIP-seq signal (Supplementary Fig. 13b, c). Components with localized signal at the respective enhancer are speculated to regulate the target genes. For the integration of ChIP-SICAP data with DepMap data (release: 21Q2), for a given target gene regulated by MIBC enhancer and occupied by FRA1 and FLI1 (according to ChIP-seq data), we identified the genes on which the respective target gene has been determined to be co-dependent (Top100 genes identified with either Crispr and/or RNAi screens ^64, 65^, positively correlated ones).

### Examination of the expression of key MIBC genes in different stages of MIBC patients

We used TCGA 2017 cohort ^8^ to check MIBC driving genes’ mRNA expression difference between different histopathological stages of the tumor samples. We grouped tumors in T2 stage (n = 132) representing the “less-aggressive” and T3 and T4 stages together (n = 277) representing “more-aggressive” and compared the mRNA expression of *FLI1* and *MAP4K4* genes between these two different groups using cBioPortal ^57^ with its “compare” option. Boxplots representing the mRNA expression change between different tumour stages were also obtained from cBioPortal ^57^.

## Results

### Enhancer landscapes differentially characterizing NMIBC and MIBC

Using the H3K27ac ChIP-seq data we generated for 2 NMIBC and 4 MIBC cell lines, we called the active regulatory elements in NMIBC (n=31911) and MIBC (n=58226) cell lines. Our analysis identified that 37% of NMIBC regulatory elements overlapped with MIBC regulatory regions (Fig. 1a, Supplementary Fig. 1). Performing an unsupervised clustering analysis on the regulatory regions determined for both NMIBC and MIBC was able to separate NMIBC and MIBC cell lines (Fig. 1b). Using the appropriate tools, we identified the active regulatory elements differentially regulated between NMIBC and MIBC. This analysis revealed 295 NMIBC-specific and 1404 MIBC-specific regulatory elements (adjusted p value =< 0.05) (Fig. 1c). Overall, we noticed that MIBC cell lines show more heterogeneous regulatory patterns compared to NMIBC, reflecting the known MIBC heterogeneity ^10, 23^. Representative examples of NMIBC-specific and MIBC-specific active regulatory landscapes are shown in Fig. 1d-1e. Overall, our differential enhancer analysis resulted in reliable NMIBC and MIBC specific regulatory regions which we used in our subsequent analysis.

### Target genes of differentially regulated enhancers

After finding the differentially regulated enhancers for NMIBC and MIBC, one critical question was to identify the target genes of these enhancers. It has been well established that enhancer-gene regulation is largely distance independent and depends on the 3D organization of the genome and interaction of the enhancers with the promoter regions of the genes ^66^. To identify the target genes of the enhancers specific for NMIBC and MIBC, we used the data present in GeneHancer database, which combines several features such as capture Hi-C, correlation of the expression of the genes with enhancer RNAs, expression of quantitative trait loci within the enhancer of interest and provides reliable enhancer-gene interactions ^52^. Using the information available in the GeneHancer database and combining it with the expression status of the genes in NMIBC and MIBC (Supplementary Fig. 2), we linked 70% of MIBC enhancers and 65% of NMIBC enhancers with their target genes (Supplementary Table 1 and 2). We identified that among the enhancers linked with a gene in our analysis, almost half of MIBC enhancers and NMIBC enhancers regulated a single target gene while almost a quarter regulated 2 genes (Fig. 2a, b). Further, we identified that the expression level of the target genes of MIBC enhancers were overall significantly higher in MIBC cell lines compared to NMIBC cell lines, and the vice versa for the expression of the genes linked with NMIBC enhancers (Supplementary Fig. 3).

We identified the target genes of MIBC-specific enhancers to be involved in epithelium migration (adjusted p value = 0.001), regulation of epithelial cell proliferation (adjusted p value = 0.00047). (Fig. 2c, d and Supplementary Table 3), which includes the genes like *TGFB2, HIF1A*, and *PRKCA* highly relevant for epithelial to mesenchymal transition ^67, 68, 69^. On the other hand, our analysis revealed that NMIBC-specific enhancers regulated the genes involved in regulation of transcription by RNA polymerase II (p adj. value = 9.1764066279671e-05) (Fig. 2e, f), and significantly enriched for transcription factors (16.45%, Fisher’s Exact Test odds ratio = 3.12, p value = 5.192e-07) defined by Lambert et al. ^70^, consisting of several FOX family and Zinc finger family members (Supplementary Table 4). Those also include *GRHL2*, which was previously identified as anti-oncogene in bladder carcinoma by regulating the expression of *ZEB1* gene thus preventing EMT both in cell lines and tissue samples ^71^. In addition, KDM5B, which is a H3K4 demethylase ^72^ and regulated by NMIBC specific enhancers, is also implicated in progression of bladder carcinoma ^73^. These results suggest that while the NMIBC-specific enhancers mainly regulate the transcription factors implicated in maintenance of a non-invasive character, MIBC-specific enhancers activate the genes involved in cell migration and proliferation such that the cells gain invasive characteristics.

### Transcription factors implicated in regulation of NMIBC and MIBC

Following the identification of the differential regulatory elements characterizing NMIBC and MIBC and their target genes, we tempted to uncover the transcription factors involved in the regulation of these elements. Performing a transcription motif finding revealed FOS family of transcription factors ^74^ to be enriched at MIBC-specific regulatory elements (Supplementary Fig. 4) while the transcription factors including p53/p63/p73 family ^75^, nuclear receptor family ^76^ were enriched at NMIBC-specific regulatory elements (Supplementary Fig. 5). To refine our transcription factors list and to determine the ones which would have differential activity in NMIBC and MIBC, we complemented our transcription factor motif analysis with the expression analysis of the respective transcription factors. We analyzed the expression of the top 10 transcription factors enriched at NMIBC and MIBC regulatory elements using the gene expression data available for the cell lines we use ^46^ or patients classified as NMIBC and MIBC ^48^. We identified that FRA1, a transcription factor belonging to FOS family and implicated in epithelial to mesenchymal transition and metastasis ^77, 78^ to be enriched at MIBC-specific regulatory elements and significantly overexpressed in both MIBC lines and MIBC patients. We also determined FLI1, a transcription factor involved in development and cancer migration and invasion ^79, 80^, as the next best candidate enriched for MIBC-specific enhancers though its expression in MIBC cell lines and patients was borderline significant (p-value = 0.1 for cell lines and p-value = 0.07 for tissue.) (Fig. 3a, b and Supplementary Fig. 6). On the other hand, transcription factors p63, a tumor suppressor with differential variant expression and involved in bladder tumorigenesis ^81^, GRHL2, a transcription factor supporting the expression of epithelial genes and deregulated in various cancers ^82^, and the PPARg, a nuclear receptor highly mutated in bladder cancer and implicated in urothelial differentiation ^83^ were significantly enriched at NMIBC-specific regulatory elements and significantly overexpressed in NMIBC lines and NMIBC-class patients (Fig. 3c, d and Supplementary Fig. 6). These results suggest that FRA1 and FLI1 might be the two critical drivers implicated in muscle-invasive characteristics of bladder cancer while the proper function and existence of p63, GRHL2 and PPARg are required to preserve non-muscle invasive properties.

**Fig. 3.**
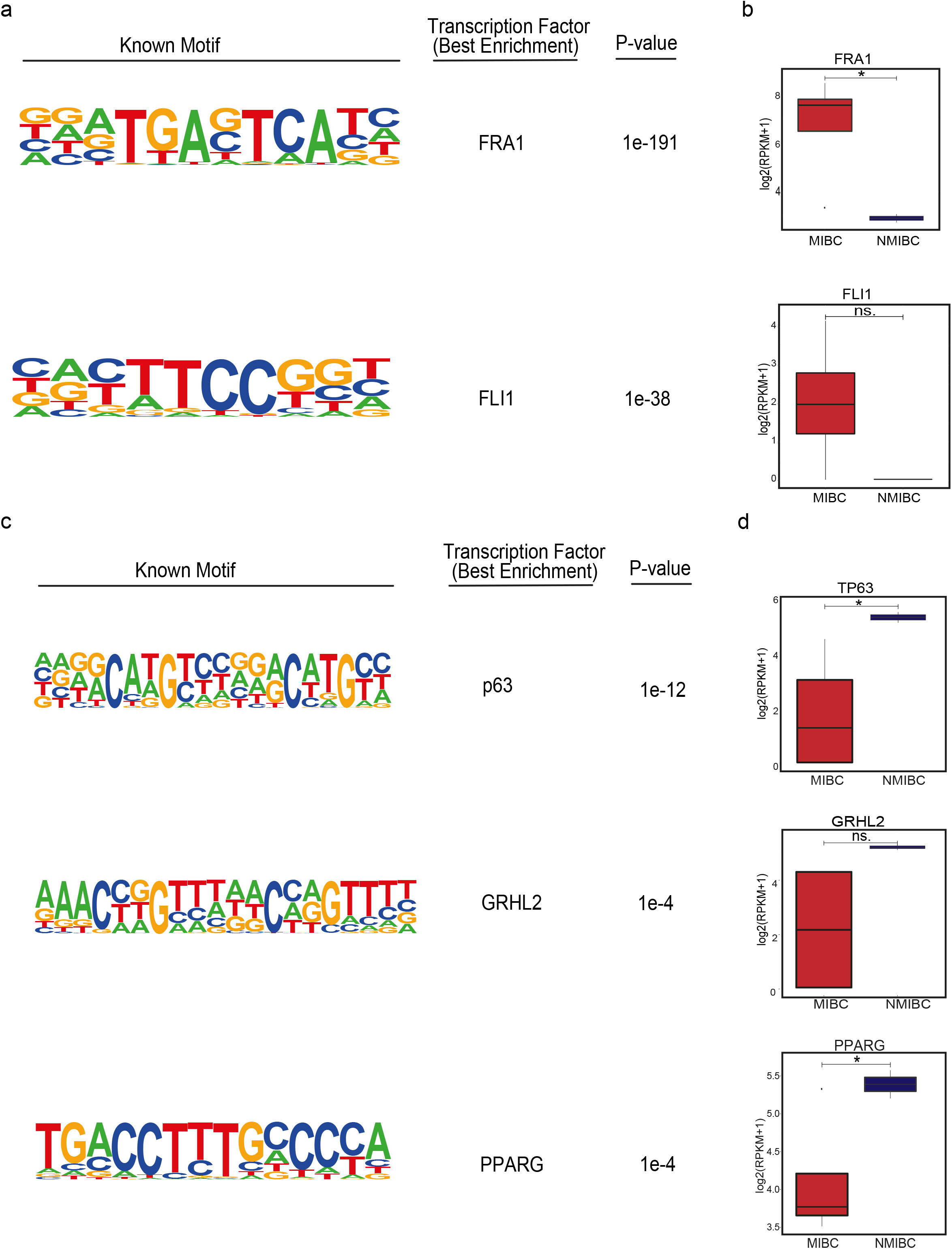
Transcription factors enriched at MIBC and NMIBC enhancers. (a, c) Top transcription factor motifs enriched at MIBC (a) and NMIBC (c) enhancers. (b, d) Boxplots showing the expression of *FRA1* and *FLI1* across the NMIBC (n=2) and MIBC cell lines (n=4) (b) and the expression of *TP63, GRHL2* and *PPARG* (d). * p value < 0.05

**Fig. 4.**
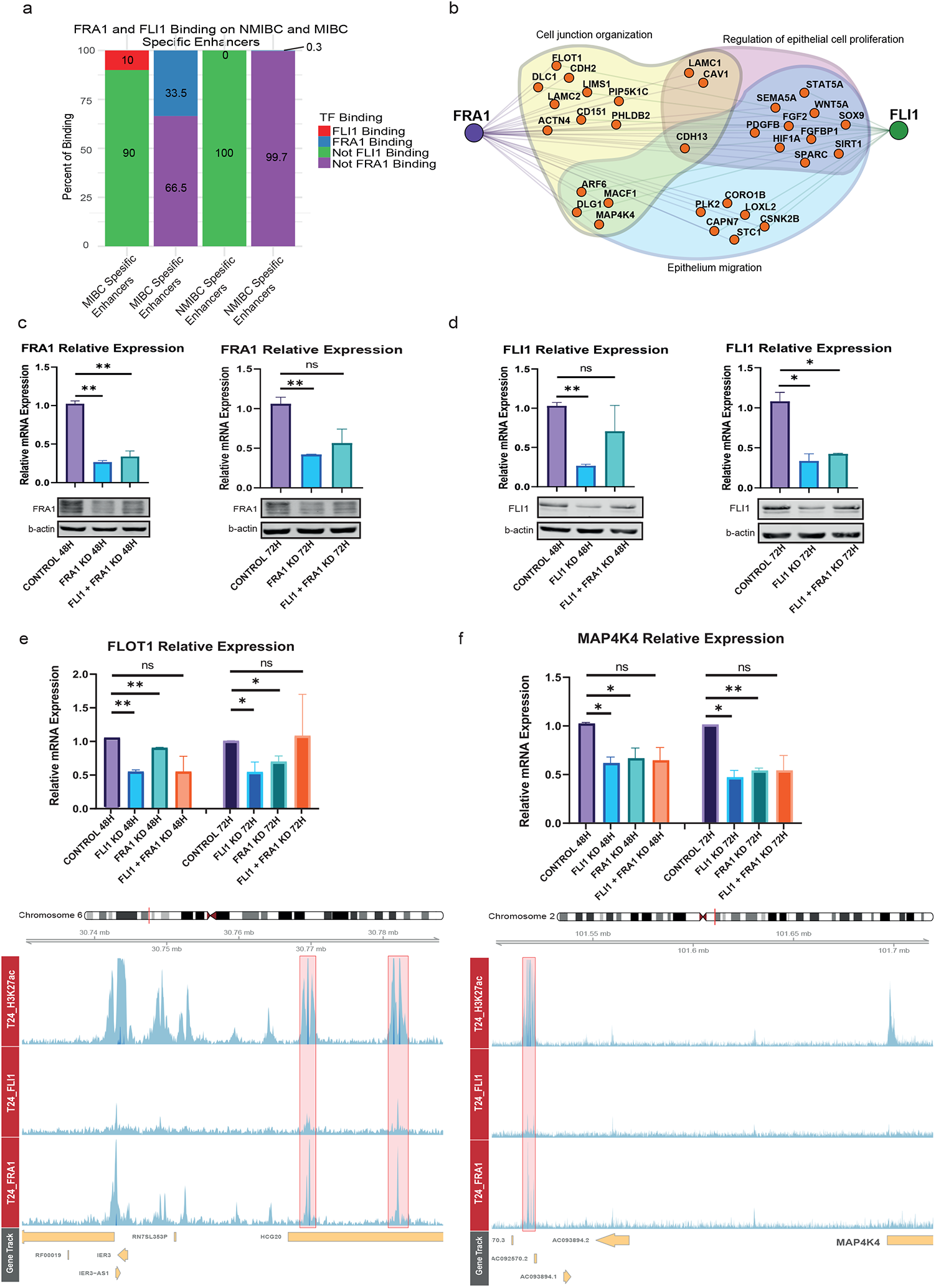
Knockdown of FRA1 and FLI1 downregulates the expression of genes involved in epithelial cell migration. (a) Barplots show the overlap rates of MIBC and NMIBC enhancers with FRA1 and FLI1 peaks. (b) Regulatory network denoting the transcription factors, FRA1 and FLI1 and their target genes involved in epithelium migration, cell junction organization and regulation of epithelial cell proliferation. For simplicity similar terms were left out of the visualized image (please see Supplementary Table 3, ‘epithelium migration’ and ‘cell junction organization’, and ‘regulation of epithelial cell proliferation’ related terms are highlighted in yellow). (c, d) Plots showing the downregulation of FRA1 (c) and FLI1 (d) at mRNA and protein level. Top panels display the relative mRNA levels, bottom panels show the Western blot images. (e, f) Barplots display the relative mRNA levels of *FLOT1* (e) and *MAP4K4* (f) (top panels). Snapshots showing H3K27ac, FRA1 and FLI1 signals at the MIBC enhancers regulating *FLOT1* and *MAP4K4* genes (bottom panels). Scale of the snapshots are adjusted to 25. Error bars show the standard deviation of two biological replicates. Each biological replicate is analyzed in three technical replicates.* p value < 0.05, ** p value < 0.01

**Fig. 5.**
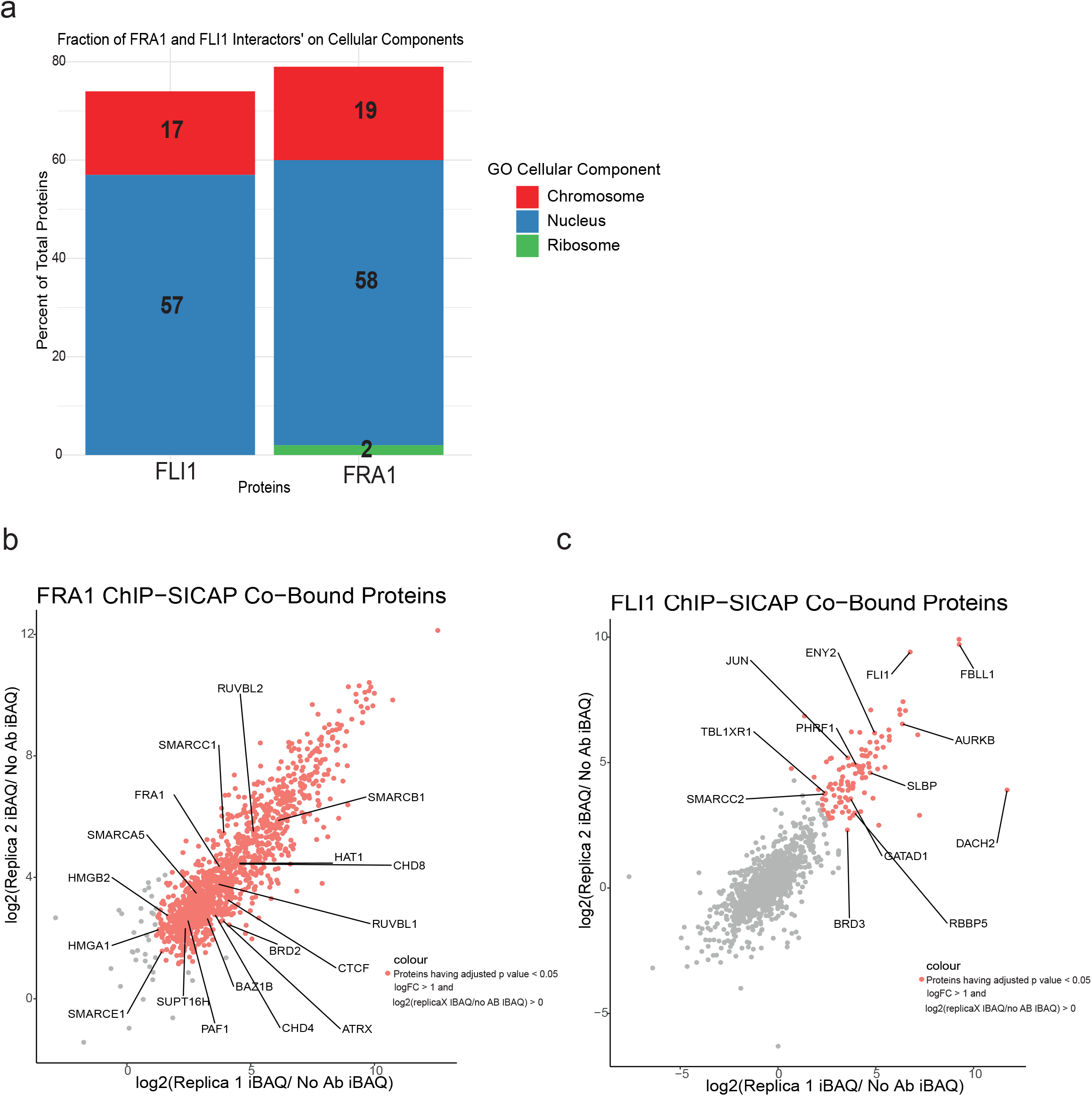
Chromatin factors interacting with FRA1 and FLI1 in MIBC. (a) Barplots show the cellular compartments FLI1 and FRA1 chromatin-bound interactors are localized. (b, c) Scatter plots visualizing the correlation between ChIP-SICAP replicates for FRA1 (b) and FLI1 (c). IBAQ values are normalized to no antibody control values.

### FRA1 and FLI1 regulate the genes involved in epithelial cell migration and cell junction organization

Muscle invasive bladder cancer patients have very bad prognosis compared to non-muscle invasive bladder cancer and have high risk of metastasis ^2^. Therefore, a thorough understanding of the gene regulation in MIBC is essential. To establish the transcriptional regulatory networks in MIBC, we performed ChIP-seq for FRA1 and FLI1 in T24 MIBC cell line. Our results showed the localization of these two transcription factors at the enhancer regions involved in the regulation of the respective genes. Overall, FLI1 peaks overlap with FRA1 peaks by 53 % (Supplementary Figure 7a). Concerning the association with the MIBC and NMIBC enhancers, FLI1 and FRA1 peaks intersected with MIBC-specific enhancers with 10% and 33.5% rates, respectively. As a comparison, for NMIBC-specific enhancers, these rates were 0% and 0.3%, respectively, suggesting the activity of FLI1 and FRA1 mainly at MIBC-specific enhancers (Fig. 4a). Overall, we identified lower number peaks for FLI1 (n=2775) compared to FRA1 (n=10513). This result might be attributed to the lower expression of FLI1 compared to FRA1 in T24 cell line (FRA1 expression: 7.6 (log2 (RPKM), FLI1 expression: 4.2 (log2 (RPKM)) ^46^. Nevertheless, both known and de novo motif finding analysis identified FRA1 and FLI1 as the best enriched motifs at FRA1 and FLI1 peaks (Supplementary Fig. 8a, b), respectively, showing the specificity of FRA1 and FLI1 ChIP-seq data.

Next, we linked FRA1 and FLI1 with the target genes of the MIBC enhancers (Fig. 4b) involved in ‘migration’ and ‘cell junction organization’, and ‘epithelial cell proliferation’ based on our results (Fig. 2c and Supplementary Table 3). We additionally discovered that FLI1 is a target gene of FLI1 itself and also of FRA1 (Supplementary Figure 7b). Further, the expression of FRA1 and FLI1 significantly correlates in bladder cancer cell lines (ρ=0.58, p value= 1.723e-3), collectively suggesting for the co-expression status of these two transcription factors and regulation of FLI1 expression by FLI1 itself and FRA1 (Supplementary Figure 7c). However, it is also worth to note that while expression and regulation of FRA1 shows a more uniform pattern among the MIBC cell lines analyzed (Figure 3a, Supplementary Figure 9a), this is more heterogeneous for FLI1, implying the differential activity of FLI1 in MIBC compared to FRA1 (Supplementary Figure 9b).

We knock-down FLI1 and FRA1 alone or in combination in the MIBC cell line T24 to determine whether the expression of the genes within this regulatory network will be changed after manipulation of FRA1 and FLI1. This strategy resulted in a highly efficient knock-down both at the transcript and protein level (Fig. 4c, d). We identified that some key genes implicated in EMT, epithelial cell migration and cell junction organization such as *MAP4K4, FLOT1, MYC, CSNK2B*, and *CD151* (Fig. 4e, f and Supplementary Fig. 10) were significantly downregulated after FRA1 and/or FLI1 knock-down. Among the downregulated genes in both FRA1 and FLI1 knockdowns, *MAP4K4* attracts special attention. We discovered that *MAP4K4* expression significantly correlates with FRA1 and FLI1 expression in primary MIBC (Supplementary Fig. 11a, b). Further, primary MIBC patients expressing higher levels of *MAP4K4* had significantly higher metastatic potential compared to the ones expressing *MAP4K4* at lower levels (Supplementary Fig. 11c, d) in two cohorts of the TCGA bladder cancer studies ^8, 9^, making *MAP4K4* is a prominent FRA1 and FLI1 regulated gene critical for metastasis of bladder cancer.

### FRA1 and FLI1 interact with the chromatin remodeler complexes, mediating the regulation of EMT related genes

Having identified the role of FRA1 and FLI1 in regulation of the genes implicated in MIBC, we employed the ChIP-SICAP assay ^29^, which enables identification of the factors chromatin bound together with the transcription factor for which ChIP experiment has been performed. Our results showed that protein partners significantly enriched (logFC > 1, adjusted p value < 0.05) for FRA1 and FLI1 interaction were mainly located in chromosome and nucleus (Fig. 5a). FRA1 interacting chromatin proteins (n=1436, adjusted p value < 0.05, logFC > 1) included several components of SWI/SNF complex, chromodomain and bromodomain containing modifiers, high mobility group proteins (Fig. 5b and Supplementary Table 5). FLI1 interacting regulators (n=103, adjusted p value < 0.05, logFC > 1) included also components of chromatin remodeler complexes such as SMARCC2, RBBP5, several transcription factors such as JUN, ENY2 and kinases such as AURKB (Fig. 5c and Supplementary Table 6). In both FRA1 and FLI1 ChIP-SICAP assays, we were able to locate FRA1 and FLI1 in the respective interactomes (Fig. 5b, c).

ChIP-SICAP assay provided the information for the interacting partners of FRA1 and FLI1. To associate these identified interaction partners with the transcriptional regulatory networks we determined (Fig. 4), we made use of the available ChIP-seq data generated for the components of SWI/SNF remodeling complex ^62^ across different epithelial cancer cell lines, and the information available in depMap portal ^64, 65^ (Supplementary Fig. 12a). Using this strategy, we were able to constitute the hypothetical regulatory hubs for the regulation of *MAP4K4* and *FLOT1* (Supplementary Fig. 12b-e). We identified the components of SWI/SNF remodeler, SMARCA4, interacting with FRA1 (ChIP-SICAP data), to localize the enhancer region regulating *MAP4K4* (Supplementary Fig. 13). Among the genes MAP4K4 show co-dependency (based on the data on depMap portal), we identified IQGAP1, FHL2, and PARP2 to be present in the chromatin associated interacting partners of FRA1 (Supplementary Fig. 12b). Extending these predictions, we determined that interacting partners of IQGAP1 (String database ^84, 85^) include many proteins which are also identified in FRA1 ChIP-SICAP (Supplementary Fig. 12c). Within the regulatory hub of *FLOT1*, we also determined components of SWI/SNF complex to localize to the enhancer region regulating *FLOT1* (Supplementary Fig. 13) and identified REXO4, CLIC1, GNL1, and FAM50A, which *FLOT1* is determined to be co-dependent on (depMap) in FRA1 chromatin-bound interactome data (Supplementary Fig. 12d). Similar to the regulation of *MAP4K4*, we were able to extend the regulatory scheme for *FLOT1* using protein-protein interaction data from STRING database ^84, 85^ (Supplementary Fig. 12e). Collectively, our integrative approach constitutes valuable hypothetical regulatory models which needs experimental validation in future studies for the genes which are potentially implicated in muscle invasion of bladder cancer.

### FRA1 and FLI1 knockdown reduces invasive power of muscle invasive bladder cancer cells

After seeing the implications of FRA1 and FLI1 knock-down on transcriptional level in T24 cells (Fig. 4), we investigated the effect of knock-down in invasive characteristics of this MIBC cell line. To test the invasion capability of T24 cells for the knock-downs, we used IC-CHIP assay, which enables the detection of invasion ability through matrigel across different microenvironment ^36, 37, 86^ in three conditions: Positive control (10% FBS), negative control (0% FBS) and muscle microenvironment (Fig. 6a). Similar to the knockdown experiments for gene expression analysis (Fig. 4), knockdown experiments were quite efficient (Supplementary Fig. 14a). To create ‘muscle microenvironment’, we used myoblast C2C12 cells differentiated into muscle with horse serum (Supplementary Fig. 14b). Quantification of invasive capacity of T24 cells showed that cells invaded towards the FBS-medium but not the FBS-free medium (Supplementary Fig. 14c). Presence of muscle cells in matrigel resulted in collective cell migration in contrast to the single cell migration into cell-free matrigel, reflected as shorter mean distances from the start line (Fig. 6b, c), in agreement with prior studies which showed metastasis to muscle is rare ^87^ and the migration of bladder cancer cells into the muscle occurs in a collective invasion mode ^88, 89, 90^.

**Fig. 6.**
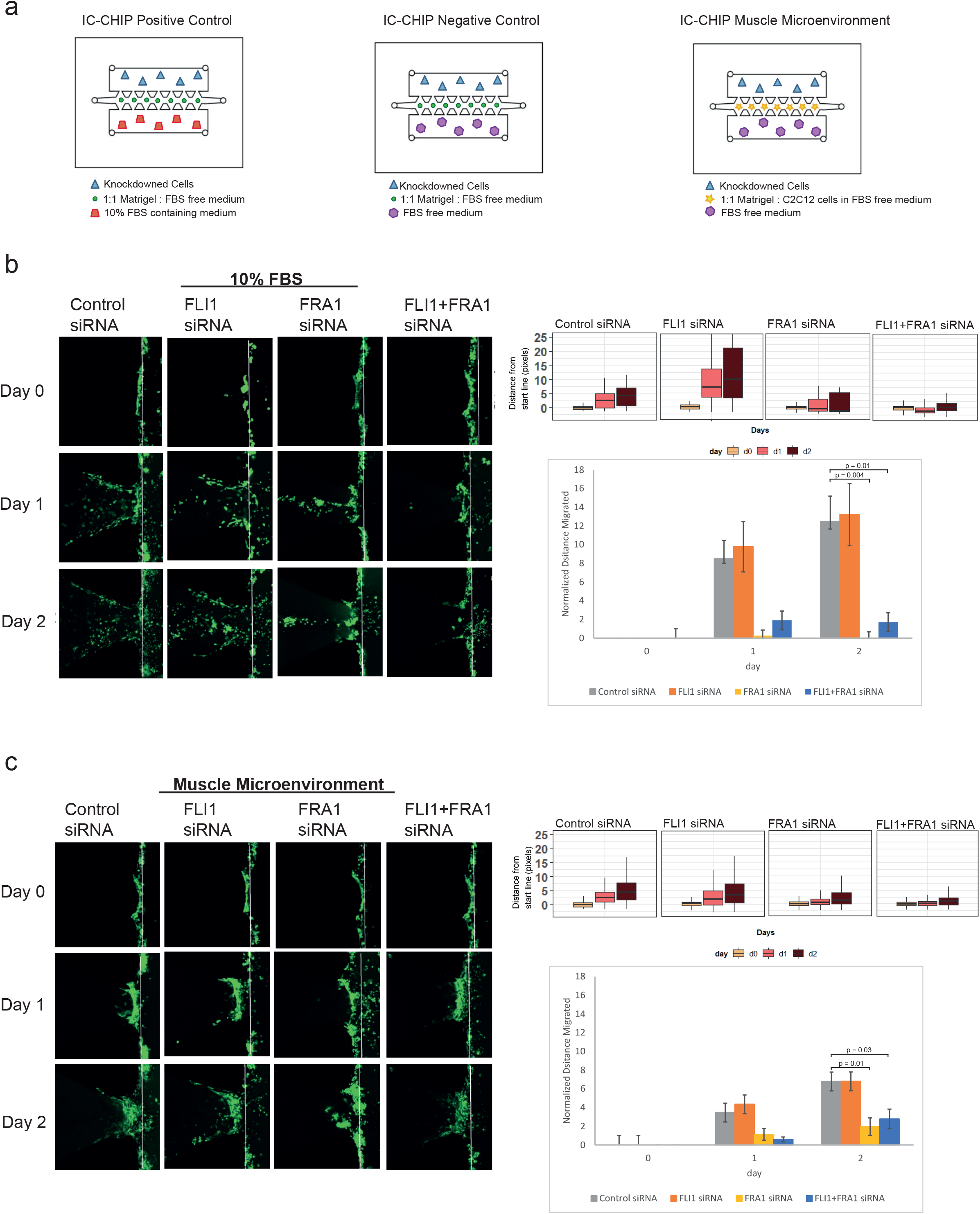
FRA1 and FLI1 knockdown results in reduction of the invasion capacity in MIBC cells. (a) Diagram showing the experimental design using IC-CHIPs. (b, c) Comparison of the invasive capacity of T24 cells into cell-free matrigel in the presence of FBS (b) and into the muscle microenvironment (c). Left panels show the representative Z-stack projection images for different conditions. Scale bar: 100µm. Boxplots display the distribution of the distance from the start line (right upper panel). Bar graphs show the mean distance values with the error bars (n=2-6) from day 0 to day 2. The data is normalized to day 0 (right lower panel). Student’s t-test (two-tailed) was used for the statistics.

Upon comparing the 10% FBS microenvironment across different conditions, compared to control siRNA, we identified significantly less invasion for FRA1 single and FRA1-FLI1 double knock-down conditions. Invasion for FRA1 single knockdown was also lower than that for FLI1 single knockdown. Invasion for control and FLI1 siRNA conditions were similar to each other. (Fig. 6b). Next, comparison of the invasion capacity towards muscle microenvironment demonstrated the role of FRA1 in invasion in the presence of muscle cells as well. Both FRA1 and FRA1-FLI1 double knock-down resulted in significantly less invasion compared to control siRNA in the presence of the muscle microenvironment (Fig. 6c). Additionally, we tested the invasion capability of another MIBC cell line, 5637 with the same FRA1 and FLI1 knock-down conditions (Supplementary Fig. 15a). In contrast to T24 cell line, 5637 cell line did not show differential invasion behavior among tested conditions in the presence of FBS (Supplementary Fig. 15b). However, similar to T24, 5637 cells showed a significant reduction in invasive capacity in muscle microenvironment upon depletion of FLI1 and also in FRA1 knock-down condition without statistical significance (Supplementary Fig. 15c), suggesting that muscle microenvironment invasion condition better mimics the in vivo conditions. Together, these results show the involvement of the FRA1 and FLI1 transcription factors in invasive capacity of bladder cancer cell lines with differential behavior in cell-free and muscle microenvironments.

We additionally overexpressed FRA1 and FLI1 transcription factors in one NMIBC cell line, RT112 to investigate potential changes on morphology and migration behavior of NMIBC cells (Supplementary Fig.16a). Overexpression of FRA1 in RT112 cells resulted in cells with enlarged cytoplasm and cell protrusions, whereas the morphological changes in FLI1 overexpressing cells were more heterogeneous with limited but detectable protrusions (Supplementary Fig. 16b). Exogenous expression of both FRA1 and FLI1 did not result in any statistically significant change in proliferation rate of RT112 cells (Supplementary Fig. 16c). However, migration rate of these cells were significantly increased upon overexpression of FRA1, as determined by wound-healing-scratch assay (Supplementary Fig. 16d).

MIBC cases show gradual muscle invasion status, starting from T2 and reaching T4 for the maximum invasion ^2^. In accordance with our findings we obtained from IC-CHIP assay, we determined significantly higher expression of *MAP4K4*, target of FRA1 and FLI1, in patients belonging to T3 and T4 groups compared to the ones which are in T2 group (Supplementary Fig. 17a). Additionally, we identified the expression of *FLI1 and FRA1* to be increased in T3-T4 stage patients as well (Supplementary Fig. 17b, c). Collectively our results highly suggest for the role of FRA1 and FLI1 in muscle invasive capability of bladder cancer.

## Discussion

In this study, we defined the active regulatory landscape of the NMIBC and MIBC cell lines using the information we gathered from H3K27ac ChIP-seq. Our in depth analysis revealed differentially regulated enhancers between NMIBC and MIBC, linked those enhancers with their target genes and identified the unique transcriptional regulatory networks implicated in NMIBC and MIBC. We further extended these analyses with transcription factor ChIP-seq and ChIP-SICAP assays, which enabled identification of the chromatin factors associated with the transcription factors involved in regulation of MIBC. Additionally, our functional studies we performed using knock-down and invasion assays proved the functionality of the transcription factors which we determined to be differentially regulated between NMIBC and MIBC.

We identified target genes of MIBC enhancers to be implicated in epithelial cell migration and cell-cell junction organization. For the NMIBC enhancers, the major target genes were related to transcriptional machinery, consisting of transcription factors, which might be critical for epithelial characteristics. In this regulatory scheme, epithelial transcription factors GRHL2, TP63 and PPARG are the main drivers constituting NMIBC regulatory network while FRA1 and FLI1, two transcription factors previously implicated in EMT in several cancers ^77, 78, 80, 91, 92^, establish the transcriptional circuitry of MIBC.

FRA1 transcription factor has been shown as motility regulator of bladder cancer cell lines RT4, RT112 and J82 ^92^. Its overexpression enhanced the mobility of RT112 cell line and its downregulation decreased the migration of J82 cell line. Moreover, FRA1 has also been shown as an important factor regulating the metastasis of squamous cell carcinoma ^91^, colorectal cancer cells ^77^. Overexpression of FLI1 has been mainly implicated in hematological malignancies ^79^. It has been shown to have a role in invasion and migration of breast cancer ^80^ and EWS-FLI1 fusion transcript has been detected in some primitive neuroectodermal tumors ^93^. Based on existing literature and our analysis, we hypothesized that FRA1 and FLI1 transcription factors can be the drivers of invasion to muscularis propria in bladder cancer. Thus, we investigated the roles of FRA1 and FLI1 in MIBC regulation in more detail. Our FRA1 and FLI1 ChIP-seq results showed the localization of these two factors at MIBC enhancers, emphasizing the role of FRA1 and FLI1 in shaping MIBC regulatory landscape. Further knock-down of FRA1 and FLI1 resulted in significantly lower expression of genes involved in epithelial cell migration. One of those genes *MAP4K4*, a kinase implicated in cancer development ^94^, has been also shown to have a role in invasiveness of bladder cancer cell lines ^95^. In addition, *MAP4K4* was one of the significantly upregulated genes in a carcinogen-induced mouse bladder cancer ^96^. It has been also shown to promote invasion in medulloblastoma cells ^97^. *FLOT1*, a member of integral membrane with a prominent role in cell morphogenesis ^98^, was another gene significantly downregulated after both FRA1 and FLI1 knock-down. This gene was identified to affect EMT transition in lung cancer ^99^, knock-down of FLOT1 in breast cancer cells resulted in decreased rates of proliferation ^100^. Further, *FLOT1* was determined to be overexpressed in bladder cancer and its expression correlated with migration and invasion characteristics of the cells ^101^.

Integration of ChIP-seq results with existing data and ChIP-SICAP suggested some previously unknown regulatory interactions, which might be critical for invasive characteristics of muscle invasive bladder cancer. Among the interaction partners of FRA1, we determined *IQGAP1, FHL2, PARP2 and CAB39* genes which MAP4K4 shows co-dependency. IQGAP1 functions in both nucleus and cytoplasm with many different roles and complex interactions, implicated in cell motility ^102^. Further, it is determined to affect cell proliferation in multiple myeloma ^103^, regulates centrosome function in breast cancer ^104^, and is involved in EMT regulation in gastric cancer ^105^. We identified CD44 and RAC1, which are identified in FRA1 chromatin-bound interactome as interactor partners of IQGAP1, further strengthening the link between FRA1 and IQGAP1. FHL2 has been previously identified to regulate the expression of p21 ^106^. PARP2 is involved in DNA repair ^107^ with high expression levels in acute myeloid leukemia ^108^. Although we could not identify a study showing the presence of CAB39 in nucleus, CAB39 protein is known with the ability of activating STK11/LKB1 ^109^ and in a preprint, it was recently shown that CAB39 protein positively regulates epithelial to mesenchymal process in bladder cancer and its knockdown decreases the migration ability of T24 and 5637 cell lines ^110^. Regarding the regulatory hub of FLOT1, localization of CLIC1 has been detected in cytoplasm, nucleus and membrane of the cells and CLIC1 has been implicated in invasion and progression of cancer ^111, 112^. FAM50A has been also shown to localize in nucleus during ameloblast differentiation ^113^. REXO4 has been shown to be implicated in drug sensitivity ^114^ and to regulate the expression of QR gene ^114^ in breast cancer. Lastly, GNL1 functions as a nucleolar GTPase and has been identified to be overexpressed in bladder cancer ^115^. Our results showed that many interacting partners of GNL1 ^84^ can be also identified among the proteins identified in FRA1 interactome, suggesting the putative role of GNL1 complex in association with FRA1 in regulation of FLOT1 gene.

We strengthened the association of FRA1 and FLI1 in bladder cancer cell’s invasion ability via employing IC-CHIP assay in different microenvironments and different knock-down conditions. By mimicking the in vivo tumor microenvironment in vitro, we were able to determine invasive power of cancer cells. Interestingly, the effect of knockdowns on invasion was influenced by the presence of muscle cells in the microenvironment, suggesting that invasion assessment should involve presence of stromal cells. What is more, with FRA1 and FRA1 and FLI1 co-knockdowns, cells showed collective rather than single cell migration, in agreement with the known role of FRA1 in cell junctions ^91^ and our findings, which showed the regulation of many genes involved in cell junction organization by FRA1 and FLI1 (Fig. 4b). Previously, FRA1 has been shown to be involved in motility of bladder cancer cells using transwell migration assays ^92^. Here, our results complement this finding via showing its role in invasion through matrigel towards serum and muscle environment, in agreement with our molecular findings. Although FLI1 knock-down has been identified to decrease migration and invasion of hepatocellular carcinoma cell lines ^116^, the role of FLI1 in invasive characteristics of bladder cancer cells has not been shown in any other study. Additionally, considering the higher expression of FRA1 and FLI1 and their targets (Supplementary Fig. 17) in bladder cancer with stages > T2, we propose that these two transcription factors can be critical players in the regulation of muscle invasive bladder cancer cell lines. Further long term invasion experiments including different cell lines and more focused inter-molecular interaction investigations can be carried out in future research; however, this is beyond the scope of this work.

Collectively, our results show the multi-faceted role of FRA1 and FLI1 transcription factors in regulation of the invasive characteristics of MIBC. We believe these results have profound implications for the specific diagnosis and treatment of bladder cancer. MAP4K4 is regarded as a novel target for several cancers ^94^, also considering that many MAP4K4 inhibitors are available, which are shown to be effective for survival of cardiomyocytes ^117^, prevention of thrombosis ^118^ and reduction of the invasion capacity of glioblastoma ^119^ and medulloblastoma ^97^, targeting MAP4K4 in bladder cancer might be a valuable strategy as well. Additionally, there are many studies investigating the effect of small molecules and inhibitors on targeting of FLI1, especially within the context of EWS-FLI1 fusion ^79^. In this regard, stratification of MIBC patients according to FLI1 expression and subsequent treatment with FLI1 inhibitors might be highly beneficial treatment strategy for MIBC patients showing high expression of FLI1. Further, a study showed the dependency of EWS/FLI1 transcription factors on bromodomain-containing proteins for the proper transcriptional regulation and treatment with JQ1, an inhibitor for bromodomain-containing proteins ^120^. In our analysis, we identified BRD3 as one of the chromatin-bound interacting partners of FLI1 (Fig. 5c). Thus, we suggest that FLI1 expression level might be a determining factor for treatment of bladder cancer patients with JQ1, which is shown to be quite effective for bladder cancer ^121^.

## Author Contributions

P.Y.G.S., G.Ö.Y. S.S. and S.E.O. designed the study. P.Y.G.S, G.Ö.Y, A.K., E.C., A.E. performed the experiments. P.Y.G.S, G.Ö.Y, A.K., A.E., H.U., G.K., D.P.O. analyzed the data. D.P.O., S.S., S.E.O. supervised the study. P.Y.G.S., G.Ö.Y., D.P.O. and S.E.O. wrote the manuscript.

## Materials & Correspondence

Requests on reagents and resources can be directed to Serap Erkek Ozhan.

## Supplementary Information for

**Supplementary Figure 1.**
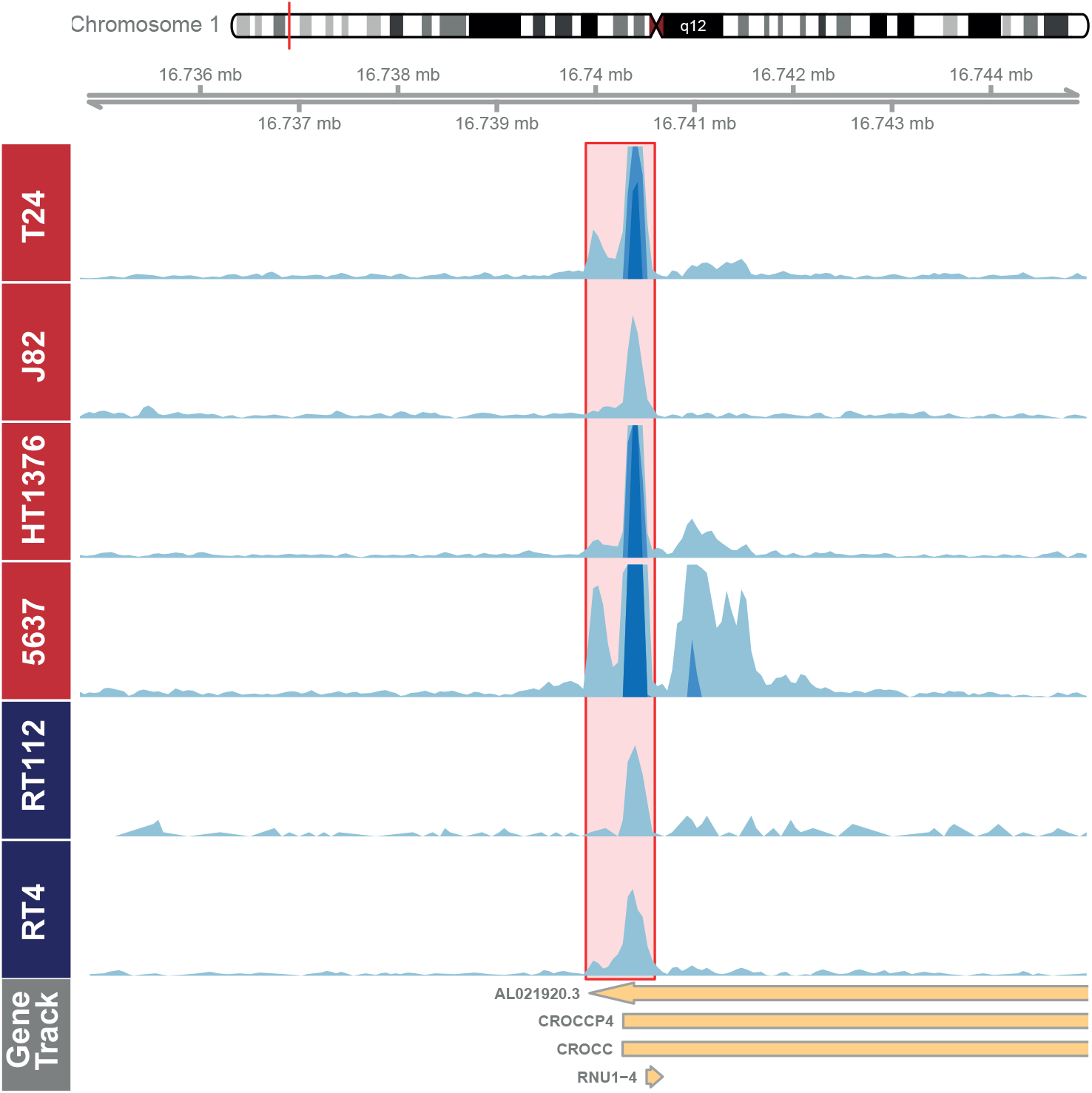
Example of a common enhancer. Snapshot visualizes H3K27ac signal at exemplary specific enhancer for NMIBC and MIBC cell lines. Scale of the snapshot is adjusted to 50.

**Supplementary Figure 2.**
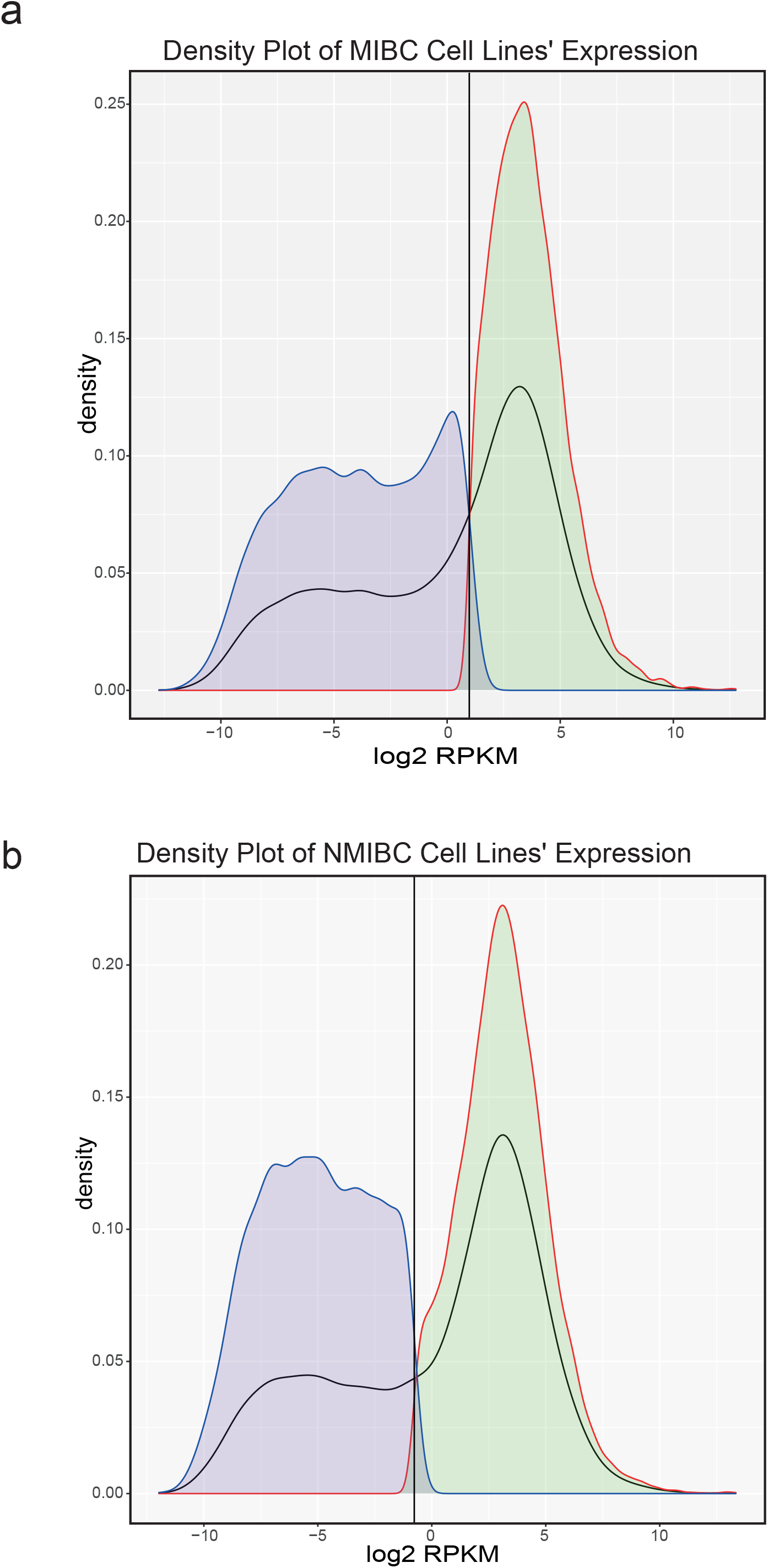
Distribution of the gene expression values in NMIBC and MIBC cell lines. (a-b) Density plots displaying the average gene expression values in MIBC (a) and NMIBC (b) cell lines. Expressed genes were shown in green and not-expressed genes are shown in purple (using “G=3” option and p-value <0.05).

**Supplementary Figure 3.**
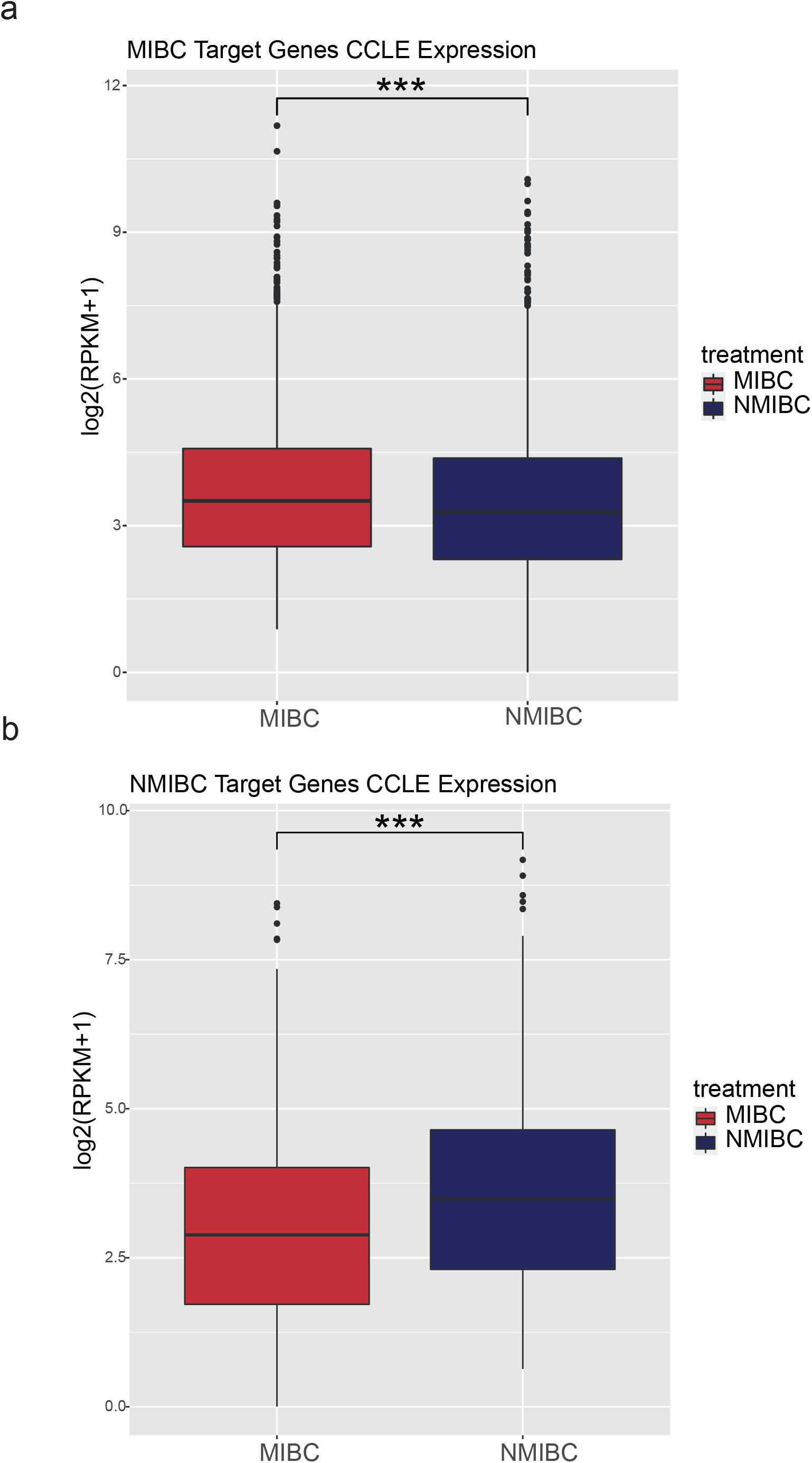
Comparison of the expression of MIBC and NMIBC target genes in cell lines. Boxplots showing the expression of MIBC (a) and NMIBC (b) target genes across the NMIBC (n=2) and MIBC cell lines (n=4). *** p value < 0.000001

**Supplementary Figure 4.**
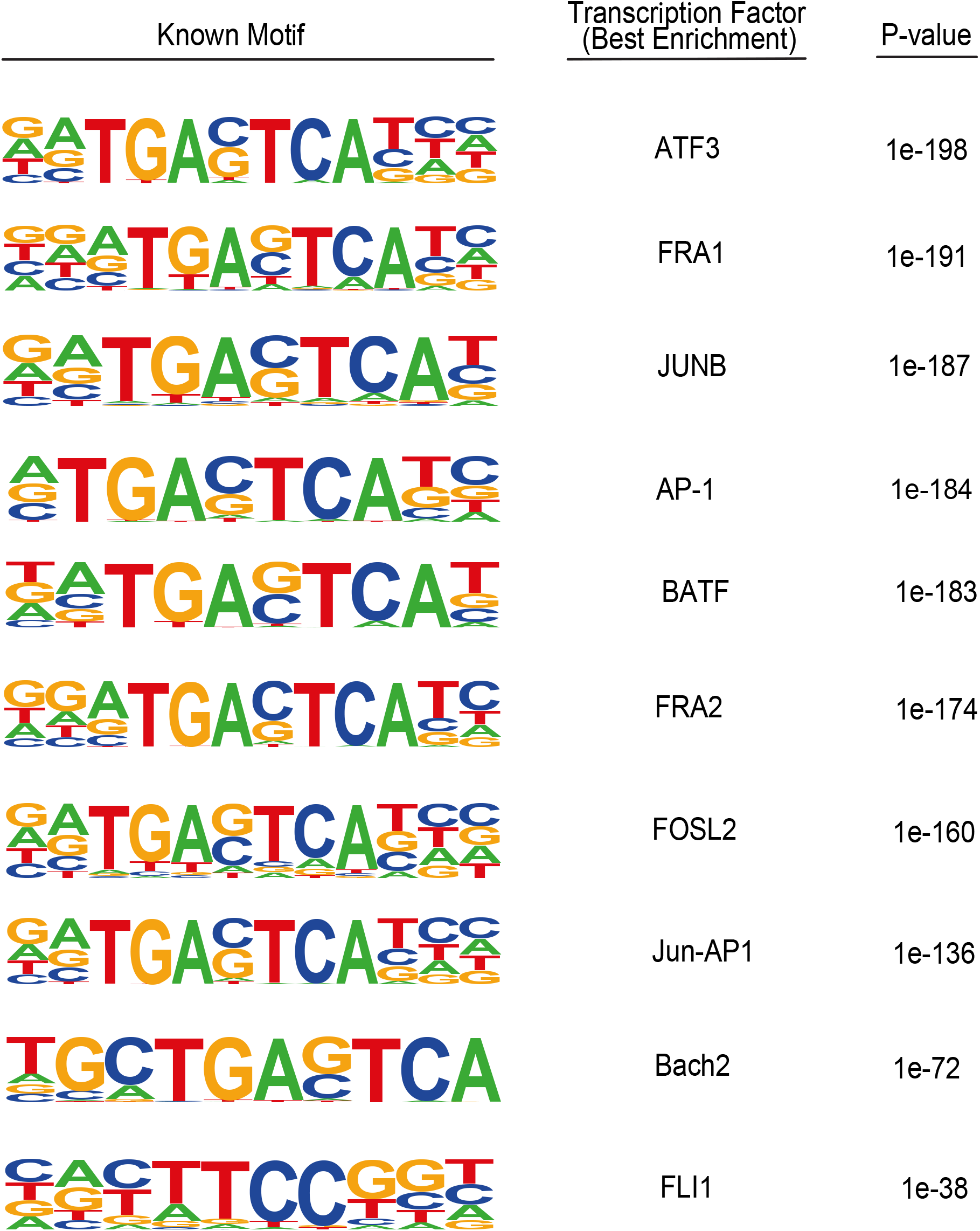
Transcription factor motif enrichment analysis in MIBC cell lines. Image shows the top 10 transcription factor motifs enriched at MIBC enhancers.

**Supplementary Figure 5.**
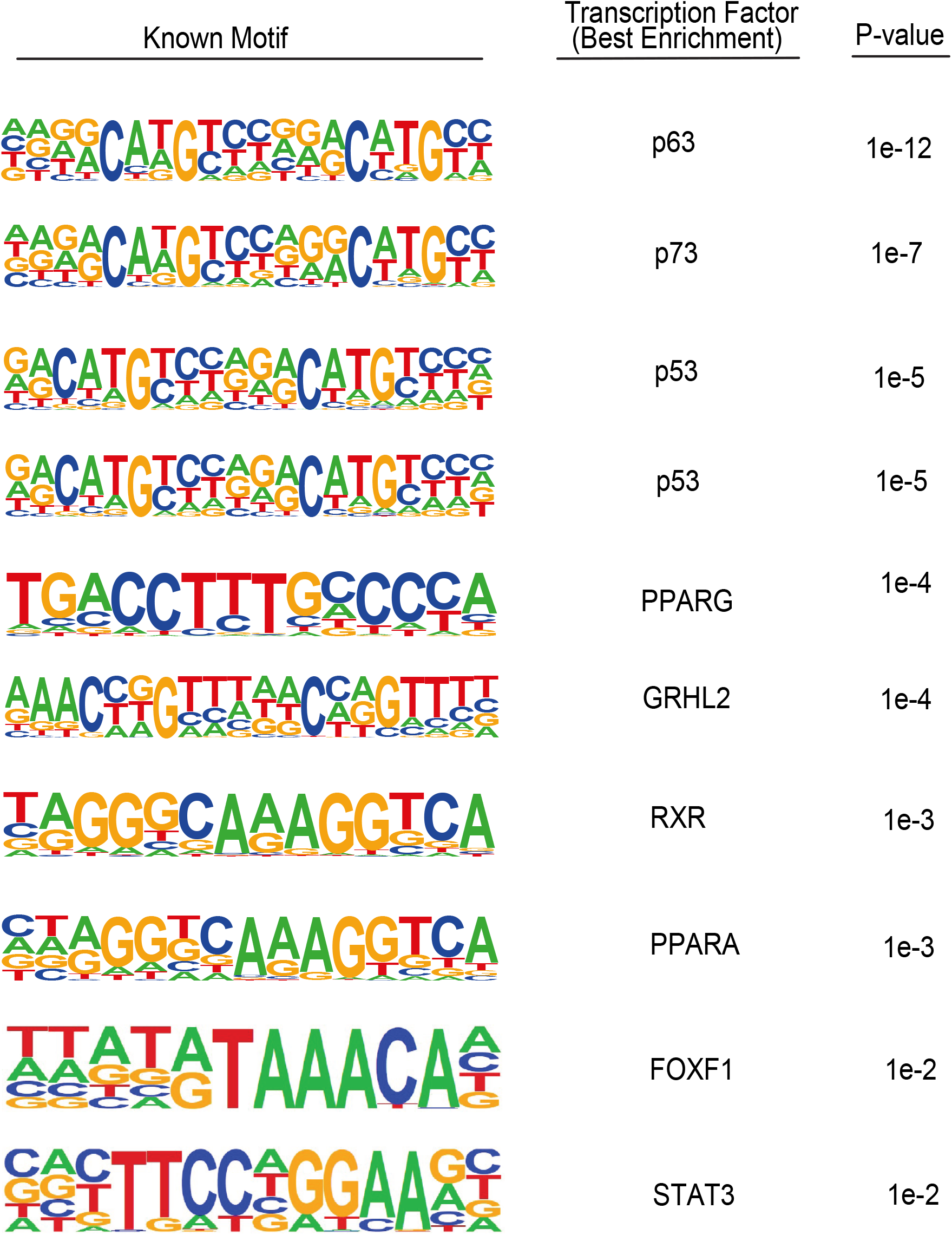
Transcription factor motif enrichment analysis in NMIBC cell lines. Image shows the top 10 transcription factor motifs enriched at NMIBC enhancers.

**Supplementary Figure 6.**
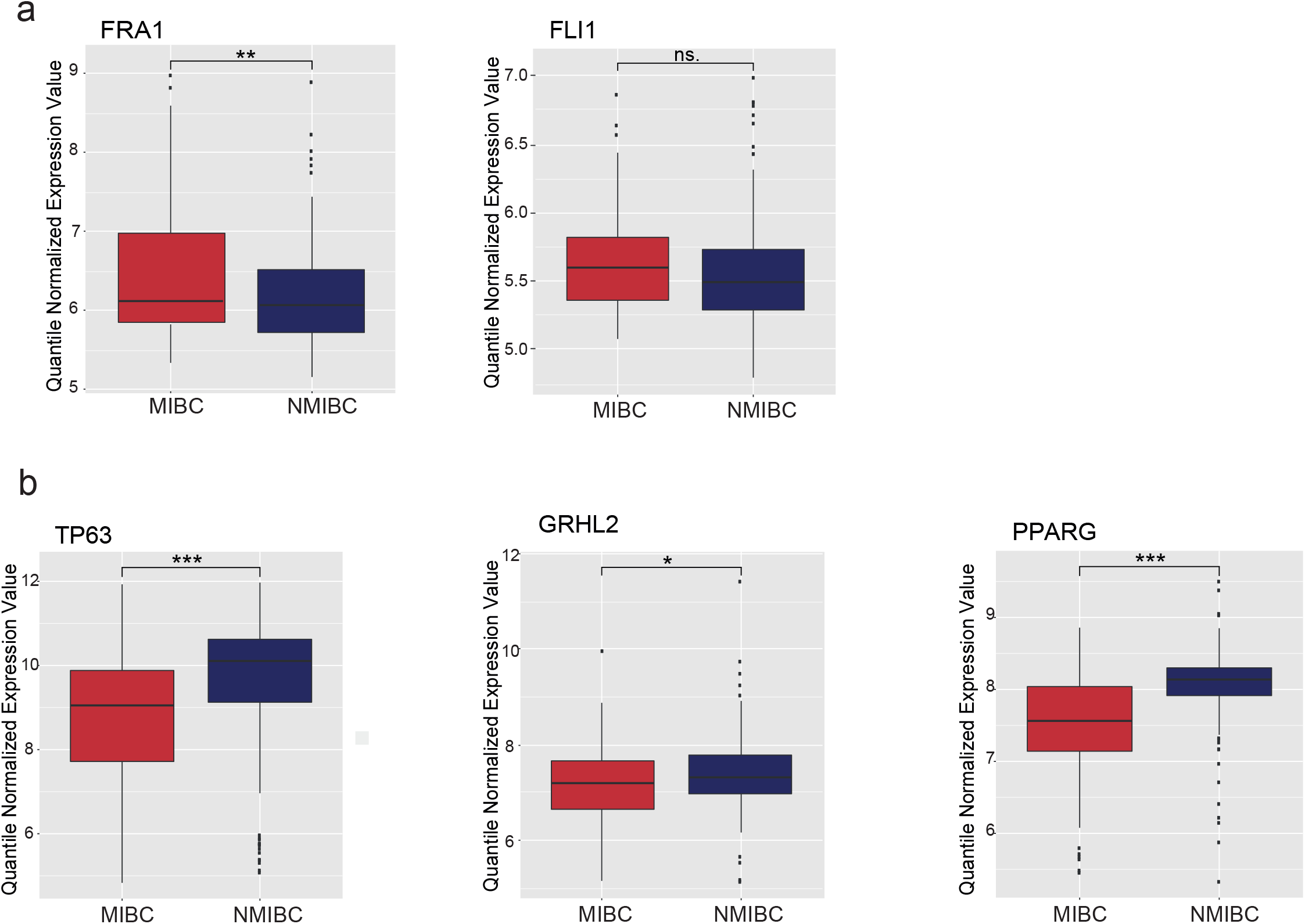
Expression level of transcription factors in primary MIBC and NMIBC tissue samples. (a-b) Boxplots display the expression level of FRA1 and FLI1 (b) and the expression of TP63, GRHL2 and PPARG across the NMIBC (n=213) and MIBC primary tissues (n=93). * p value < 0.05, ** p value < 0.01, *** p value < 0.0001, ns = not significant

**Supplementary Figure 7.**
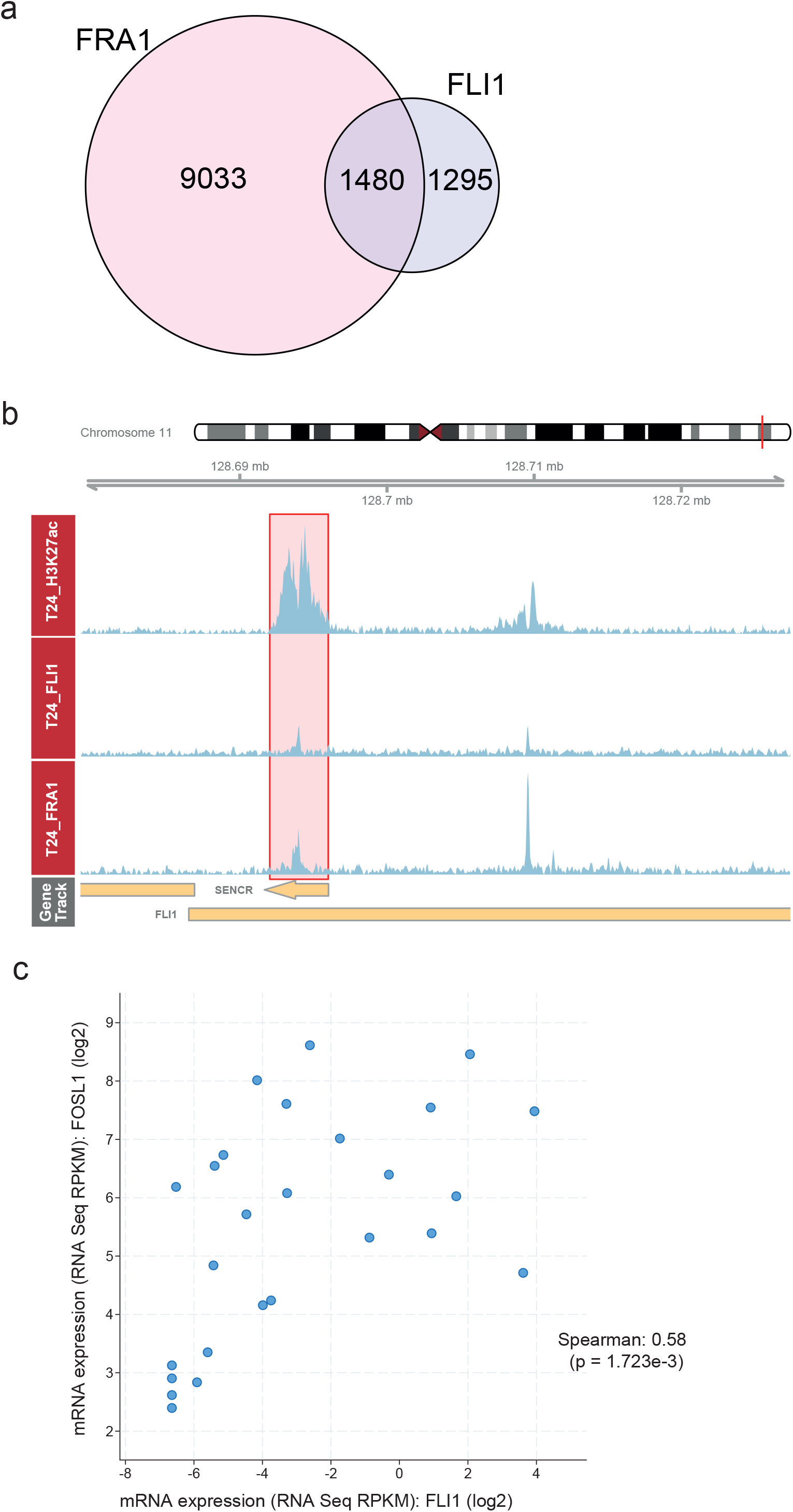
Co-regulation of FRA1 and FLI1. (a) Venn diagram showing the overlap between FRA1 and FLI1 peaks in T24 cell line. (b) Snapshot showing H3K27ac, FRA1 and FLI1 signal in T24 cell line at *FLI1* locus. Region specified by red rectangle shows the enhancer regulating FLI1.Scale of the snapshot is adjusted to 25. (c) Scatter plot showing co-expression status for FLI1 and FRA1.

**Supplementary Figure 8.**
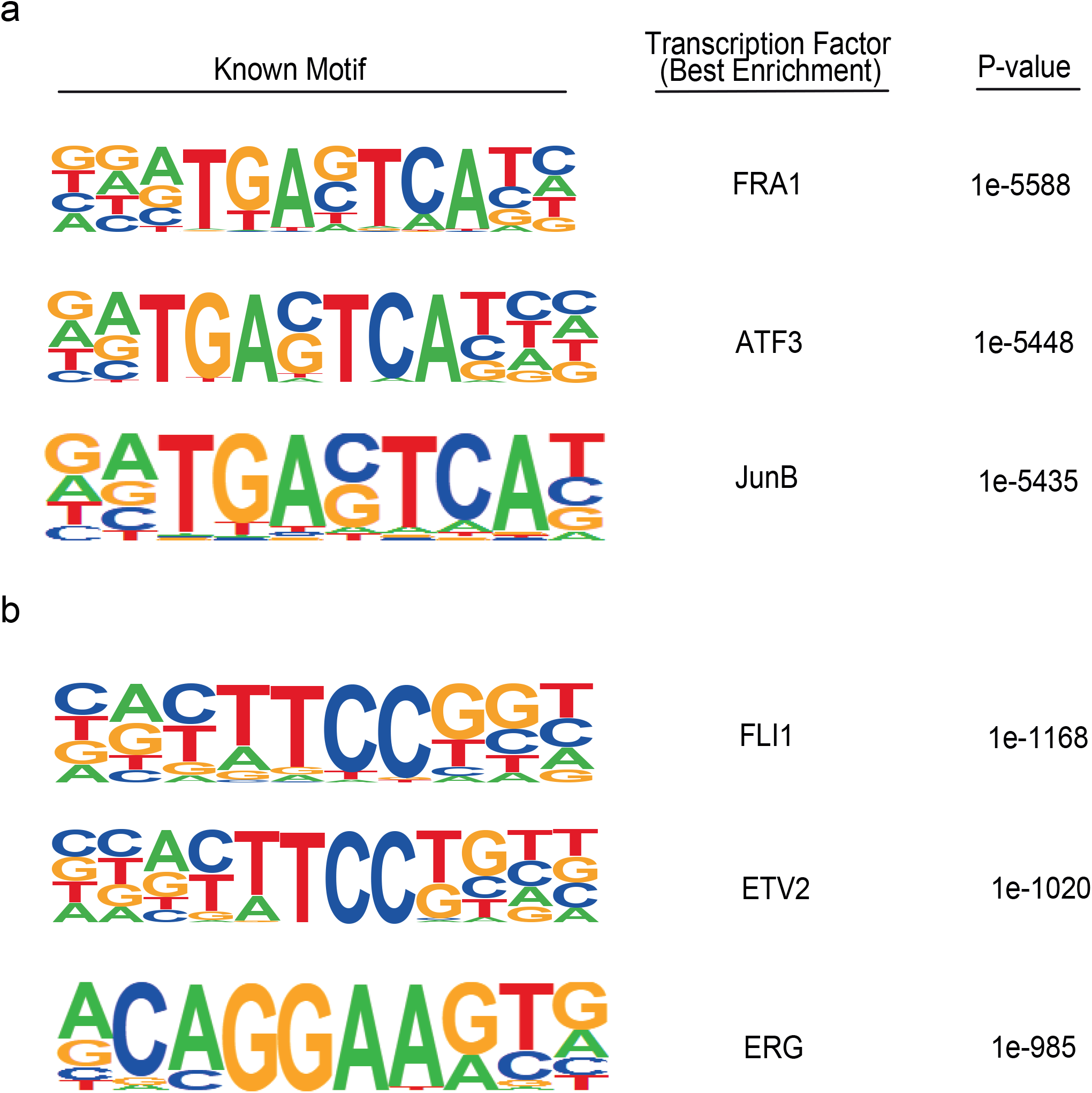
Motif enrichment analysis at FRA1 and FLI1 ChIP-seq peaks. (a-b)Top three transcription factor motifs enriched for FRA1 (a) and FLI1 (b) ChIP-seq data.

**Supplementary Figure 9.**
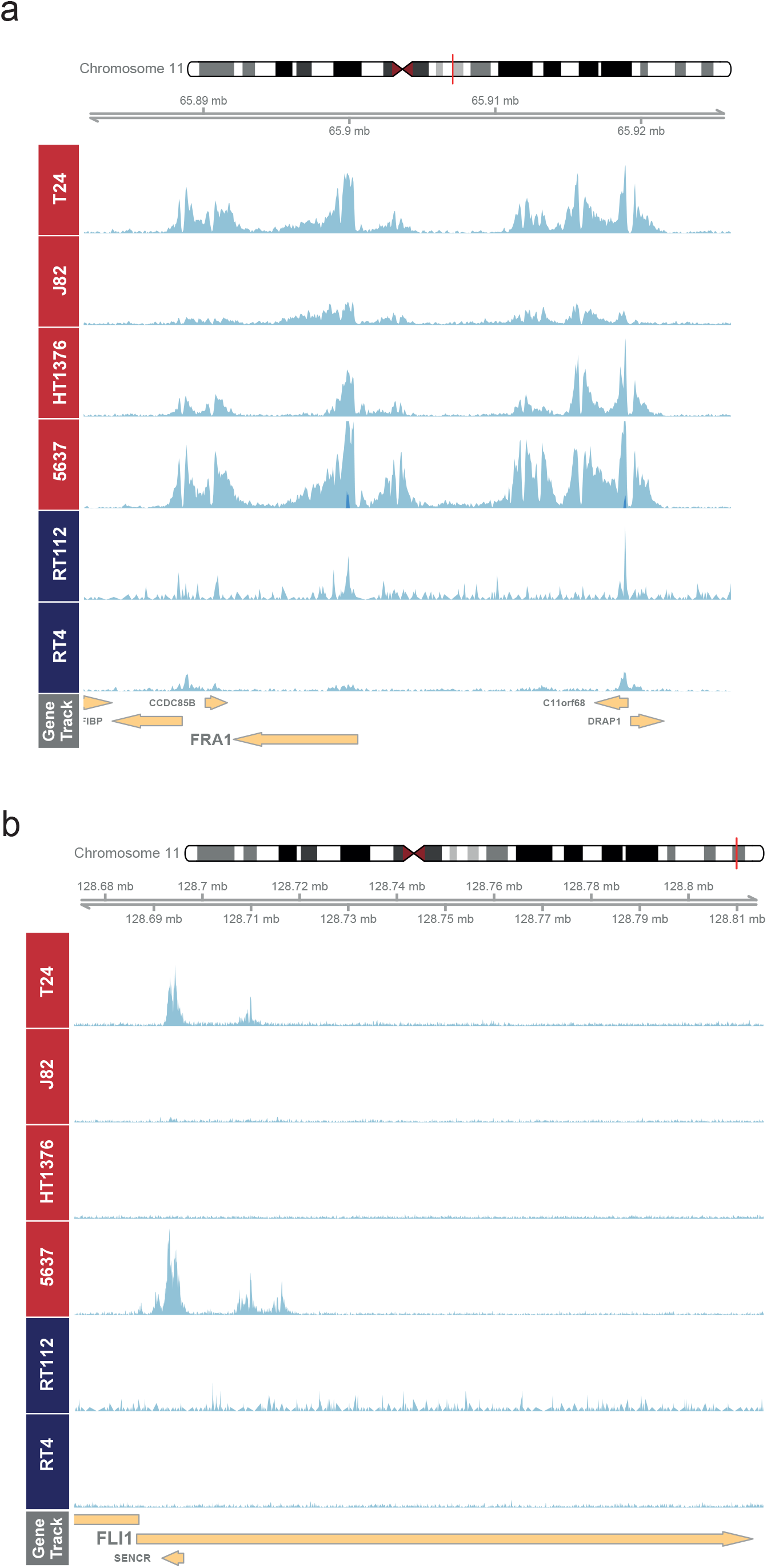
H3K27ac occupancy profiles of FRA1 and FLI1. Snapshots showing H3K27ac signal in MIBC and NMIBC cell lines at *FRA1* (a) and *FLI1* (b) loci. Scales of the snapshots are adjusted to 50 and 35, respectively.

**Supplementary Figure 10.**
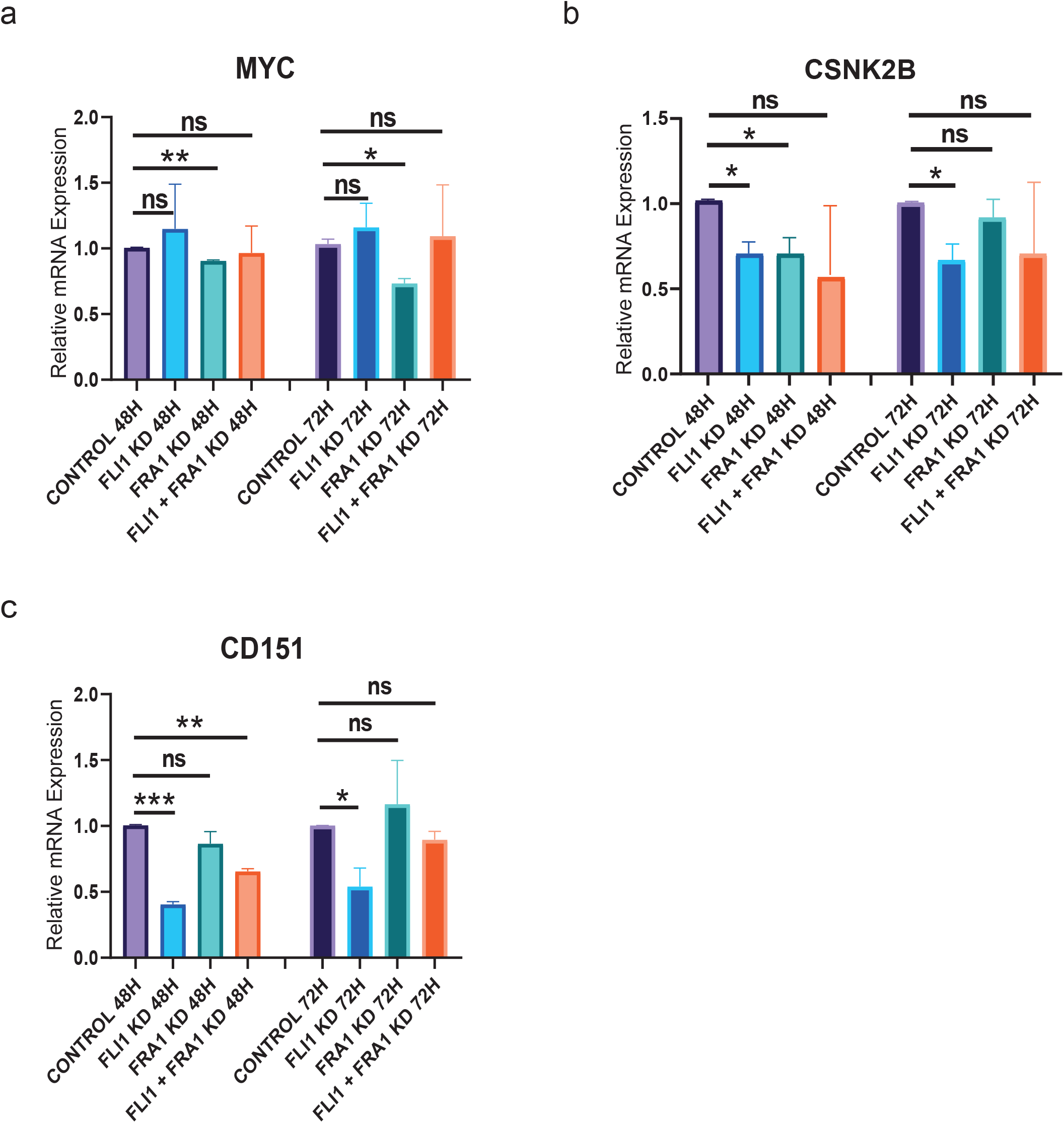
Expression level of EMT-related genes after FRA1 and FLI1 knockdown. (a-c) Barplots display the relative mRNA levels of MYC (a), CSNK2B (b) and CD151 (c). Error bars show the standard deviation of two biological replicates. Each biological replicate is analyzed in three technical replicates.* p value < 0.05, ** p value < 0.01, *** p value < 0.001, ns = not significant.

**Supplementary Figure 11.**
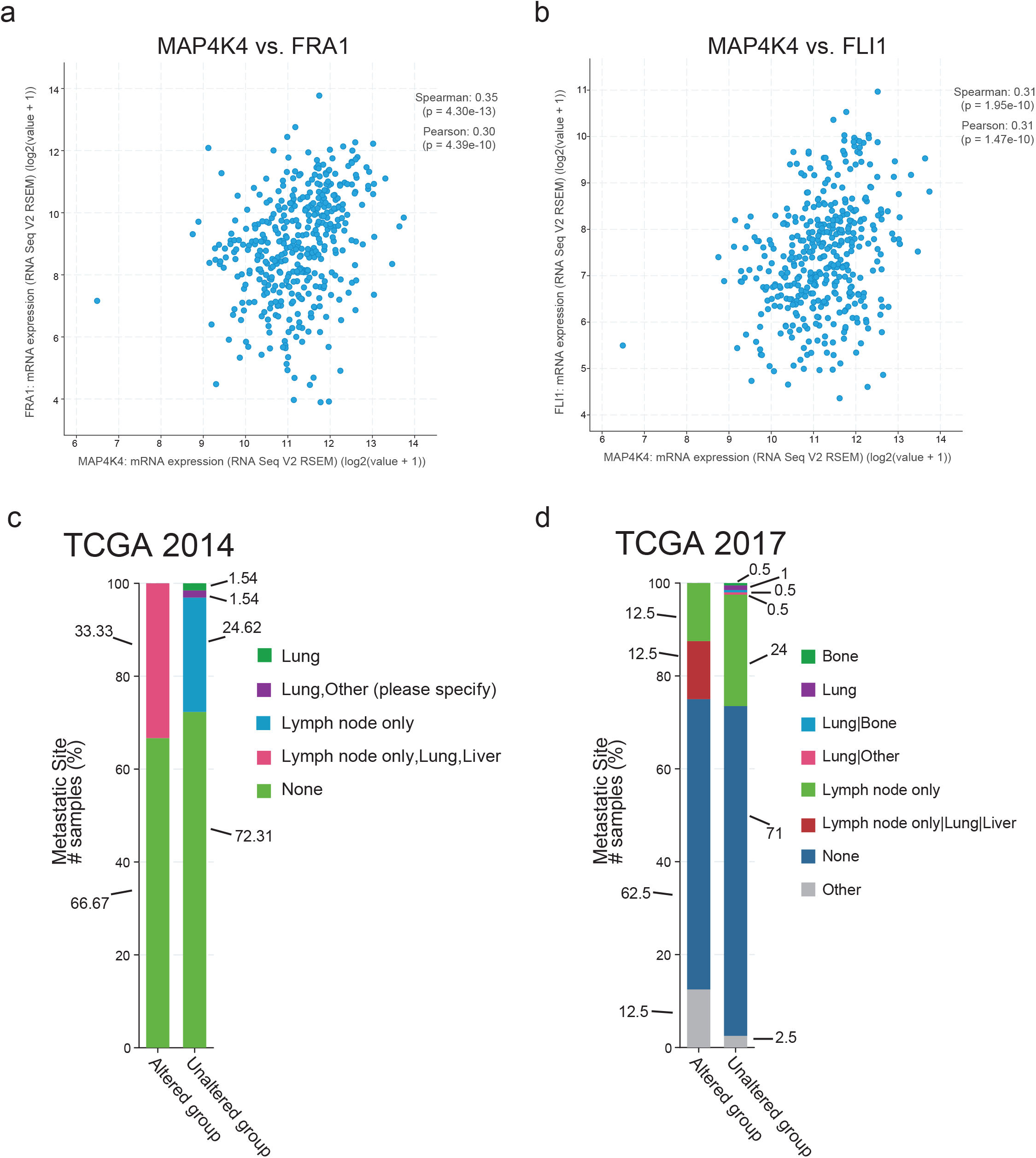
Association of MAP4K4 expression with metastatic potential of MIBC. (a-b) Scatter plots display the correlation between the expression of FRA1 (a) and FLI1 (b) with MAP4K4 expression in primary tissue. (c-d) The samples with high MAP4K4 expression is determined in cbioportal database (83) (see Methods). Barplots show the distribution of metastatic sites in the patients with high expression and not high MAP4K4 expression in two different cohorts,TCGA 2014 (c) (p value = 1.586e-4, q value = 5.868e-3) and TCGA 2017 (d) (p value = 1.878e-4, q value = 4.508e-3).

**Supplementary Figure 12.**
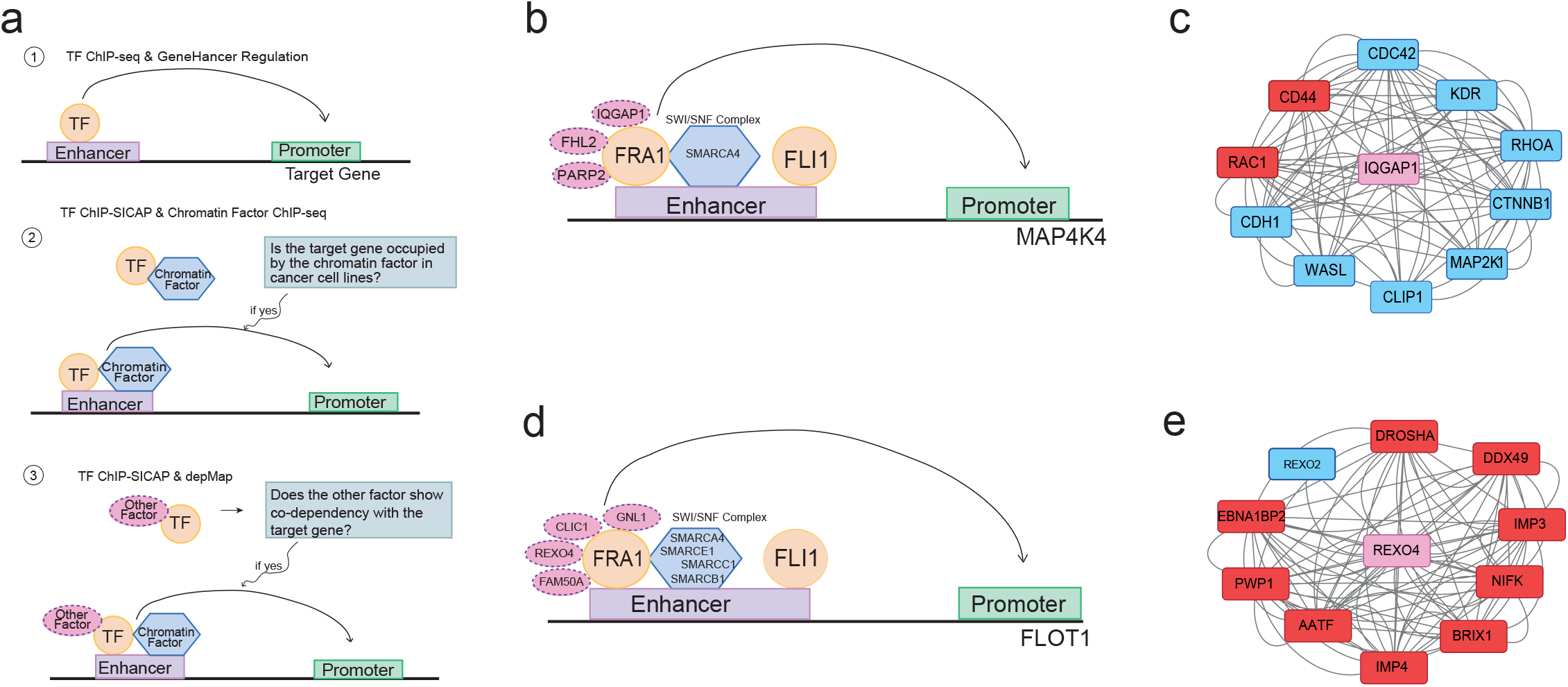
Regulatory hubs implicated in regulation of EMT-related genes. (a) Schematic showing the workflow for the constitution of the regulatory hubs. (b, d) Regulatory hubs for the regulation of MAP4K4 (b) and FLOT1 (d). (c, e) String protein-protein interaction networks for IQGAP1 (c) and REXO4 (e). Proteins which can be also identified in FRA1 chromatin-bound interactome (Supplementary Table 5) are shown in red.

**Supplementary Figure 13.**
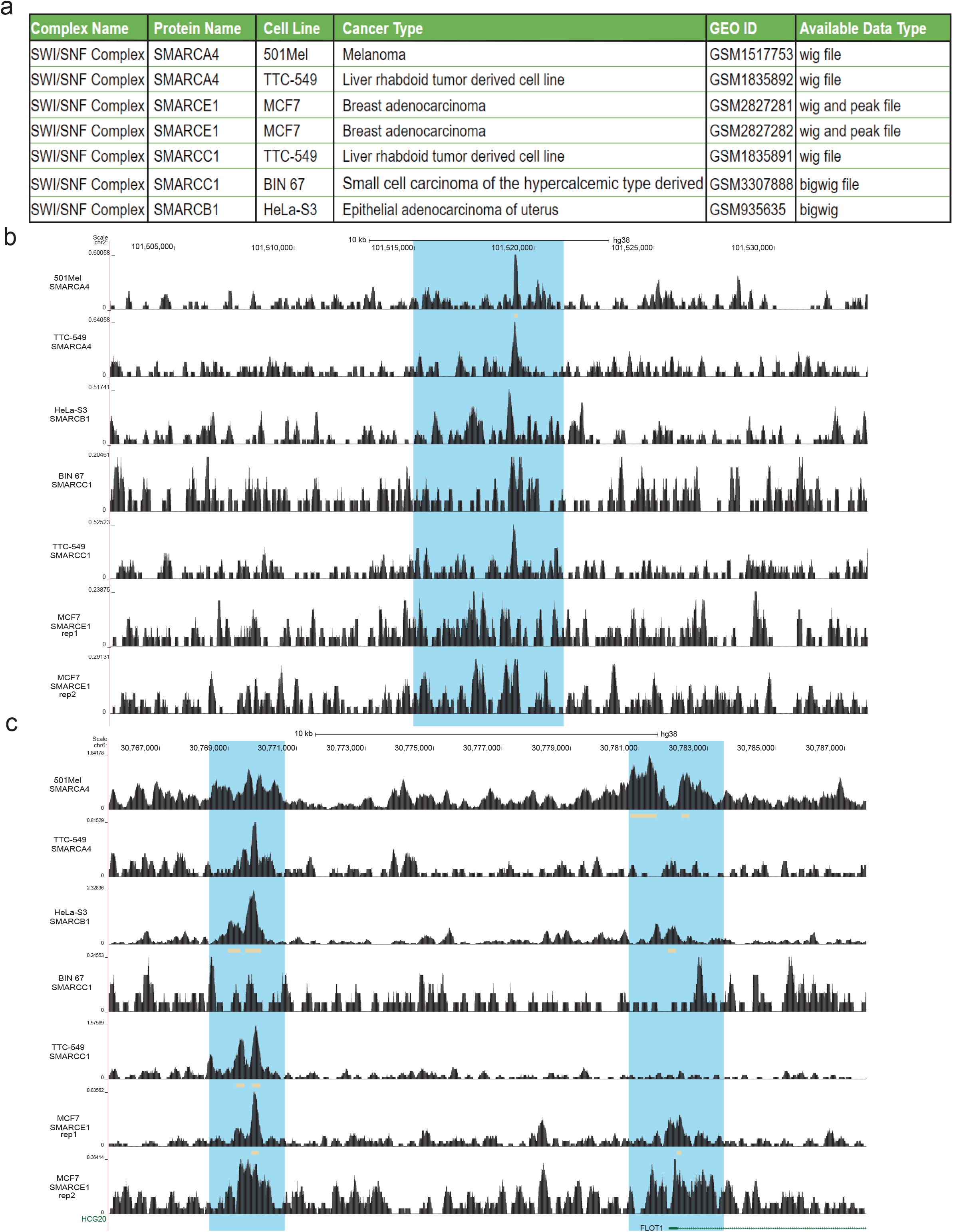
ChIP-seq signal of SWI/SNF complex components at MIBC enhancers regulating MAP4K4 and FLOT1. (a) Summary of the used datasets. (b-c) Snapshot visualizations from UCSC Genome Browser (84), showing the localization of the SWI/SNF complex components according to data from (a) at enhancers regulating MAP4K4 (b) and FLOT1 (c). Orange rectangles below the signal track show the peak region for the respective signal. Blue highlights depict the respective enhancer regions. For FLOT1 gene, two enhancer regions involved in the regulation of this gene are shown.

**Supplementary Figure 14.**
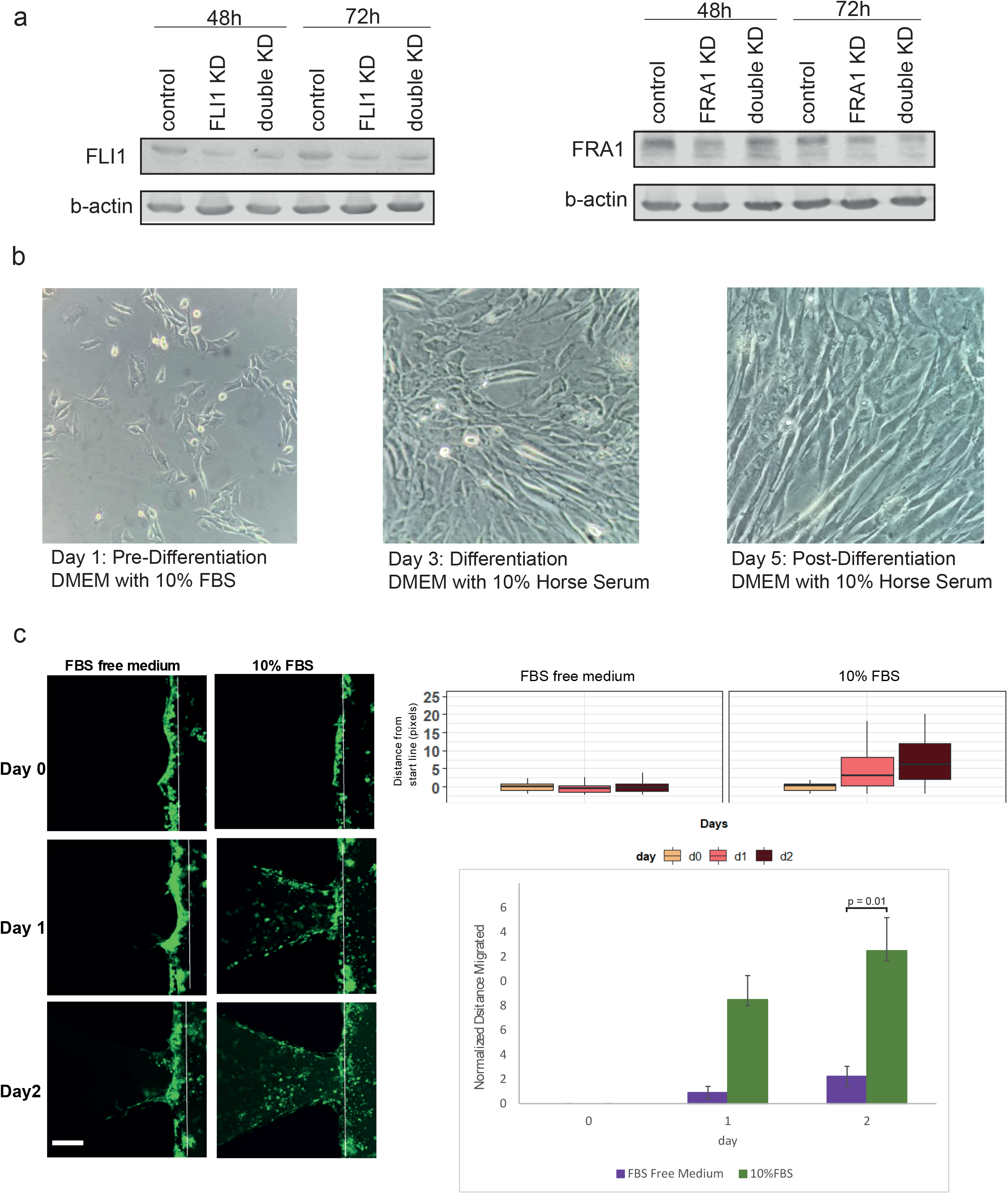
Knockdown efficiency of FLI1 and FRA1 in T24 cells used in IC-CHIP and differentiation of C2C12 cells and IC-CHIP assay in the presence and absence of FBS. (a)Western blot images show the FLI1 and FRA1 levels after FLI1 and FRA1 knock-down. The images show the differentiation of C2C12 myoblast cells to muscle for 4 days in the presence of 10% horse serum. (b) Comparison of the invasive capacity of T24 cells in the presence of 0% FBS and 10% FBS. (c) Left panels show the representative Z-stack images for different conditions. Scale bar: 100µm. Boxplots display the distribution of the distance from the start line (right upper panel). Bar plots show the distance values from day 0 to day 2. In the graphs, the data is normalized to day 0 (right lower panel).

**Supplementary Figure 15.**
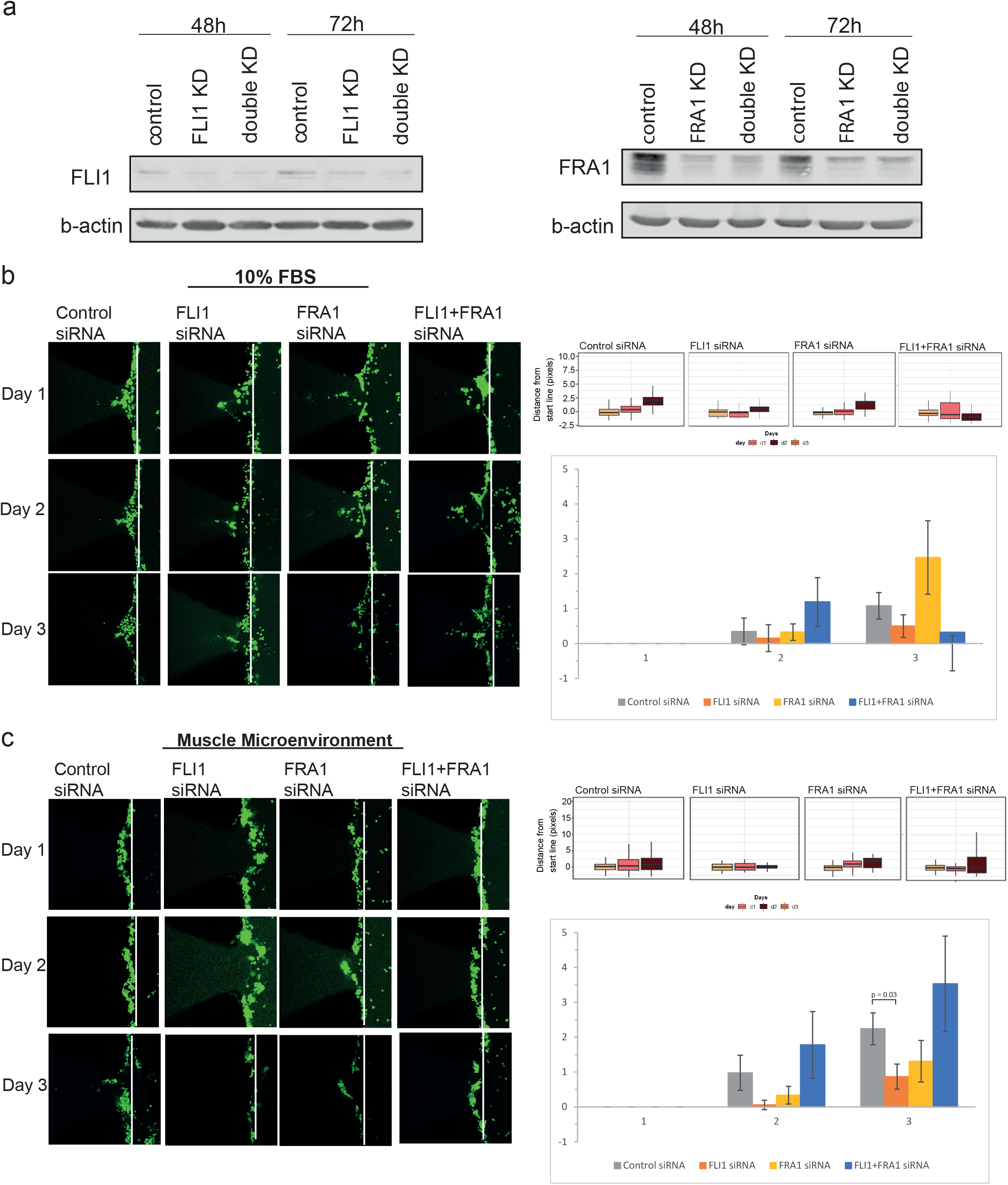
Knockdown efficiency of FLI1 and FRA1 in 5637 cells and comparison of the invasive capacity of 5637 cells into cell-free matrigel in the presence of FBS and muscle microenvironment. (a) Western blot images show the FLI1 and FRA1 levels after FLI1 and FRA1 knock-down. (b-c) Comparison of the invasive capacity of 5637 cells into cell-free matrigel in the presence of FBS (b) and into the muscle microenvironment (c). Left panels show the representative Z-stack projection images for different conditions. Scale bar: 100µm. Boxplots display the distribution of the distance from the start line (right upper panel). Bar plots show the mean distance values with the error bars (n=2-6) from day 1 to day 3. The data is normalized to day 1 (right lower panel). Student’s t-test (two-tailed) was used for the statistics.

**Supplementary Figure 16.**
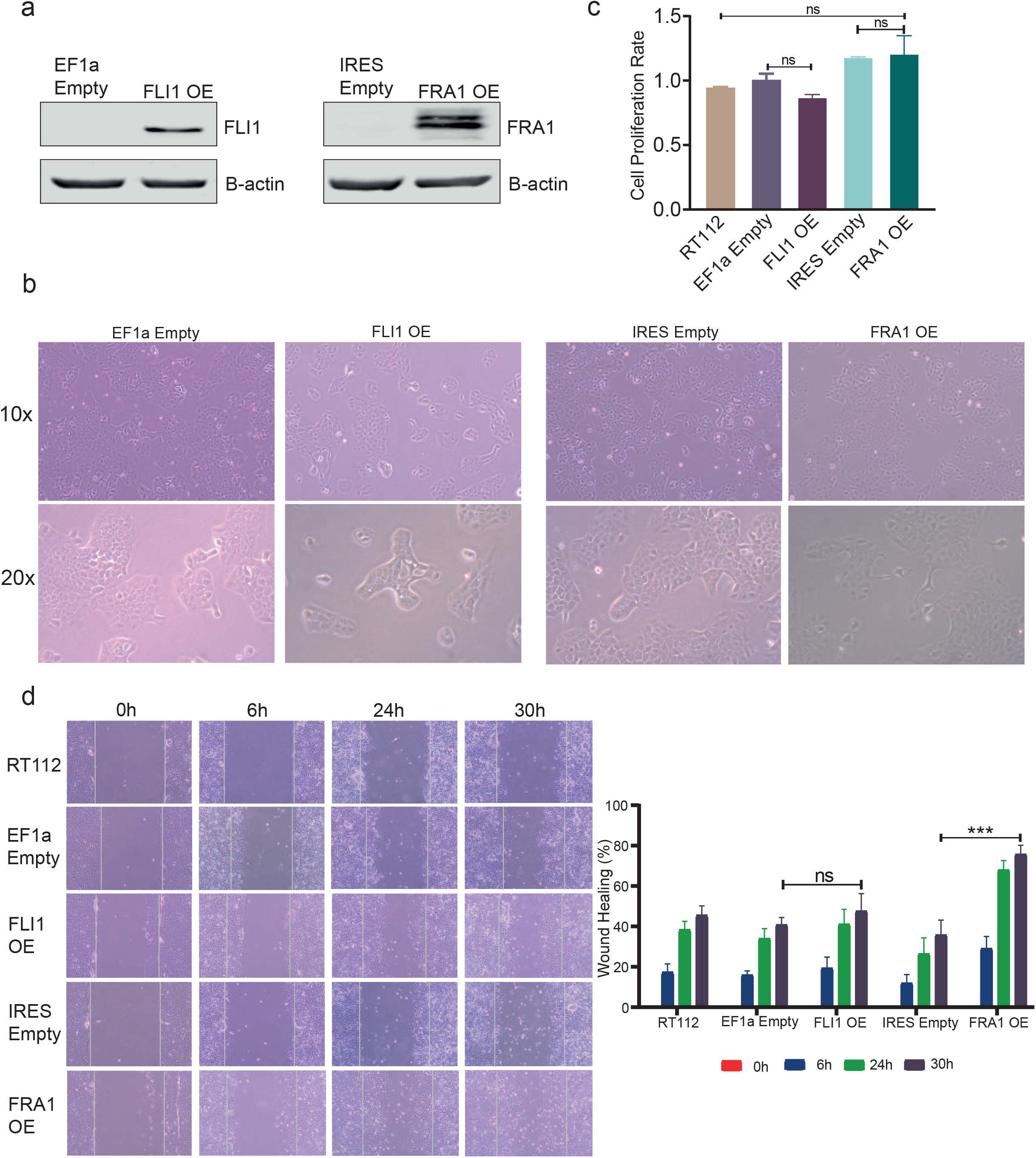
Overexpression of FRA1 and FLI1 transcription factors in RT112 cell line. (a) Western blot images showing FLI1 and FRA1 protein levels for control vector and FRA1 or FLI1 overexpressed RT112 cell line. (b) Microscope images show the morphological changes between control and FRA1 or FLI1 overexpressed cells, images were taken with 10x and 20x magnification. (c) MTT cell proliferation assay results, error bars show the standard deviation of three technical replicates. (d) Wound healing scratch assay results. Left panel shows cell images taken for different time points. Images were taken with 10x magnification. The graph in right panel shows the quantification of wound healing assay results, error bars show the standard deviation of at least 5 independent measurements. *** p value < 0.0005

**Supplementary Figure 17.**
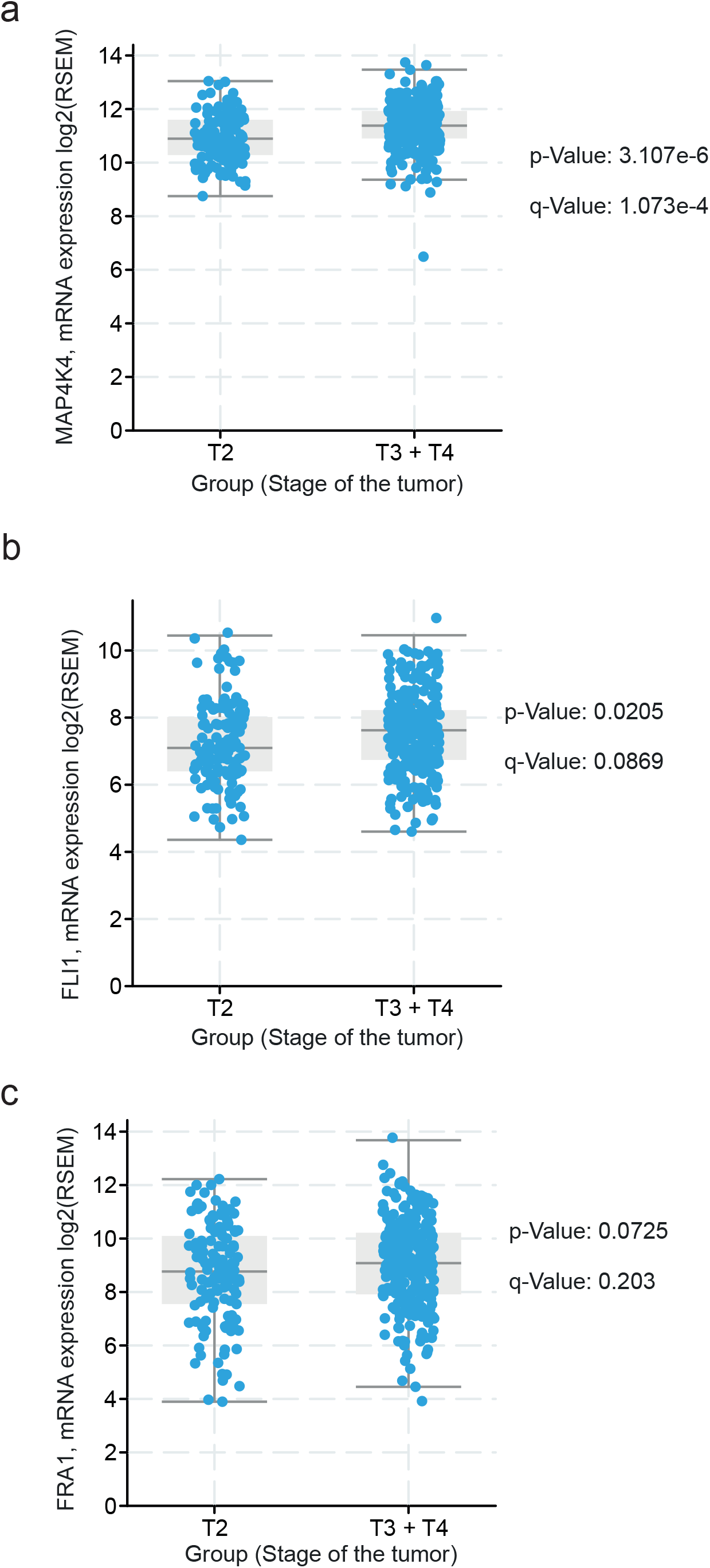
Expression of key MIBC genes in T2 vs T3-T4 stage MIBC patients. (a-c) Boxplots show the expression of MAP4K4 (a), FLI1 (b) and FRA1 (c) in T2 stage and T3-T4 stage MIBC patients.

**Supplementary Table 1**. MIBC enhancers and their target genes.

**Supplementary Table 2**. NMIBC enhancers and their target genes.

**Supplementary Table 3**. Complete results of ClusterProfiler GO Term analysis for the target genes of MIBC enhancers.

**Supplementary Table 4**. List of transcription factors identified in target genes of NMIBC enhancers.

**Supplementary Table 5**. List of proteins identified to be interacting with FRA1 in T24 cell line according to ChIP-SICAP results.

**Supplementary Table 6**. List of proteins identified to be interacting with FLI1 in T24 cell line according to ChIP-SICAP results.

**Supplementary Table 7**. Primers used in RT-qPCR experiments.

